# A portable orthogonal replication system enables continuous gene evolution near the biological speed limit

**DOI:** 10.64898/2026.02.20.706958

**Authors:** Rongzhen Tian, Fabian B. H. Rehm, Matthew Kenneth, Kiarash Jamali, Petar S. Zhotev, Kim C. Liu, Jason W. Chin

## Abstract

Orthogonal DNA replication systems uncouple the mutagenesis of target genes from host viability, enabling target gene hypermutation beyond the genomic critical error threshold and thereby unlocking access to greater sequence space for accelerated evolution. Here we introduce a series of upgrades to the *E. coli* orthogonal replication system, EcORep. We develop strategies to efficiently establish, engineer, and transform orthogonal replicons. We develop and utilize replicon-REXER to establish a 77 kb replicon, the largest orthogonal replicon reported to date. Directed evolution of the orthogonal DNA polymerase yielded variants with mutation rates of ∼10^-4^ substitutions per base per generation and best-in-class mutational spectra. These polymerases are three orders of magnitude more mutagenic than the first-generation EcORep system, enable mutagenesis at one million times the genomic levels, and straddle the evolutionary critical error threshold for the mutation of genes tested. Using the highly mutagenic EcORep system, we rapidly evolve an ethanol assimilation pathway for increased performance. Furthermore, we find that the three components sufficient to drive the minimal EcORep system enable O-replication systems to be established in other Gram-negative bacteria. Thus, we establish VinORep in *Vibrio natriegens*. VinORep combines O-replicon mutation, around the limit for molecular evolution of genes, with the fastest growing organism, to realize gene evolution approaching the biological speed limit. We exemplify the utility of this advance through the rapid evolution of new function – via the accumulation of tens of mutations, and selection – in 16 hours.

## Main

The natural evolution of new gene functions is a slow process, resulting from the accumulation of mutations in a population coupled to the selection for advantageous phenotypes. These mutations occur infrequently as organisms have evolved mechanisms to ensure their genomes are replicated at high-fidelity, thereby limiting the occurrence of mutations that could be detrimental to reproduction or survival.

Directed evolution enables the accelerated diversification and selection of genes for pre-defined phenotypes. Classical methodologies rely on *in vitro* mutagenesis steps followed by introduction of the mutagenized genetic material into an appropriate host for phenotypic selection ^1^. Generally, multiple cycles of mutagenesis and selection are necessary to obtain variants with the desirable level of fitness. Although this strategy has seen much success, the exploration of gene fitness landscapes is heavily constrained by the inherent restrictions imposed on evolutionary depth and scale by the *in vitro* diversification and transformation steps. Continuous evolution methodologies aim to sidestep these limitations via the continuous *in vivo* hypermutation of defined genes undergoing selection ^2,3^. In these approaches, the evolutionary speed depends on the rate and bias of the mutagenesis, the growth rate of the host, gene copy number, and the choice of selective pressure.

Early approaches for continuous evolution involved inserting target genes into viral genomes and infecting mutagenic cells ^4–6^. Selection for phenotypes of interest is then achieved by coupling the desired phenotype to the capacity to produce viral progeny, thereby amplifying and propagating active gene variants. These approaches, however, are limited to evolving small genes that are readily packaged into viral particles. Moreover, coupling a desired activity to viral propagation is challenging and often requires substantial case-by-case optimization.

Other approaches rely on the targeted localization of a mutagenic moiety, such as a deaminase or a mutagenic polymerase, to a defined gene ^7–12^. However, these approaches are often restricted by their mutagenic window, the types of mutations that can be accessed, and their degree of orthogonality. Where mutations can occur at elevated levels outside of the target genes, genes associated with the selection for a given phenotype may mutate and lead to selection escape. By contrast, orthogonal replication systems, where an orthogonal DNA polymerase (O-DNAP) exclusively replicates an orthogonal episome, enable robust continuous evolution experiments ^13–16^.

Essentially all existing orthogonal replication systems rely on DNAPs that replicate linear replicons via a protein-primed replication mechanism ^13–17^. The first such system, termed OrthoRep, was established by exploiting natural linear plasmids from yeast and has enabled a range of continuous evolution experiments via engineered mutagenic DNAP variants ^18–20^. More recently, an orthogonal replication system was established in *E. coli*, the workhorse of synthetic biology, by repurposing replicative components from the PRD1 bacteriophage ^14^. *E. coli* grows more rapidly and to higher cell densities than *S. cerevisiae*, key parameters dictating evolutionary speed and scale. However, the initial EcORep system relied on engineered DNAPs with mutation rates that were only modestly higher than the genomic mutation rates (2 x 10^-7^ substitutions per base, per generation). Recent work reported using a six-component system (T7 DNA polymerase, T7 helicase-primase fusion protein, host factor thioredoxin, T7 single-stranded DNA binding protein, T7 lysozyme-T7 RNA polymerase fusion protein, and T7 origin of replication) to specifically replicate a circular plasmid in *E. coli*. A mutation rate of 1.7 x 10^-5^ was reported for the plasmid ^21^.

Here, we report the development of the second-generation EcORep system. We developed highly efficient approaches to engineer orthogonal replicons and used them to generate a replicon 77 kb in length, the longest sequence maintained by an orthogonal replication system to date. Moreover, we show that replicons can be extracted and efficiently transformed into cells expressing only two EcORep components – the terminal protein and the O-DNAP. We used directed evolution to generate DNAP variants with mutation rates that are orders of magnitude higher than the initial EcORep system and exhibit best-in-class mutational bias. These O-DNAPs span the critical error threshold for 2 kb of genes encoded in the O-replicon. We employed the new highly mutagenic EcORep system to rapidly evolve the performance of a multi-gene ethanol assimilation pathway.

Next, we demonstrated that the three components that define the minimal EcORep system in *E. coli* are sufficient to establish orthogonal replication systems in other Gram-negative bacteria of biotechnological significance, *Vibrio natriegens* and *Pseudomonas putida*. By coupling the rapid generation time of the fastest growing organism, *V. natriegens*, with the rapid O-replicon mutagenesis facilitated by highly mutagenic O-DNAPs, we realized a highly accelerated continuous evolution system that surpasses existing mutation rates per day by at least an order of magnitude. This system accumulated more than 10 mutations in a target gene within 16 h and enabled selection for a new phenotype in a single day.

### Efficient establishment and engineering of EcORep O-replicons

Establishing and maintaining the EcORep linear replicon (O-replicon), harboring genes of interest, relied on the controlled expression of key replication components from the EcORep synthetic replication operon. The genes of interest within the linear replicon are flanked by inverted terminal repeats (ITRs), which function as origins of replication.

The EcORep synthetic replication operon consists of genes encoding for a terminal protein (TP) which primes replication from the replicon ends, an orthogonal DNA polymerase (O-DNAP) which specifically replicates the replicon and does not interfere with genome replication, and single-stranded DNA binding protein (SSB). The system was commonly established by electroporation of the O-replicon DNA (commonly generated as PCR amplicons) into cells that also express the PRD1 double-stranded DNA binding protein (DSB) and the Gam protein from lambda red phage.

Expanding on our previous work ^14^, we developed multiple key aspects of EcORep. Initially, we found that shortening the period of induction of Gam, SSB, and DSB expression when preparing electrocompetent cells improved replicon establishment efficiencies from PCR amplicons. The optimized process yielded ∼10^4^ transformants for short replicons (1.2 and 2.5 kb in length, 2.5 μg of purified orthogonal replicon PCR product with), with all colonies obtained harboring the desired O-replicon (Supplementary Figs. 1 and 2). Longer replicons, however, could only be established at a low efficiency and yielded a large proportion of false-positive colonies (Supplementary Fig. 2).

Next, we found that replicons could be efficiently engineered *in vivo* using the λ Red recombination system. We initially inserted an ampicillin resistance gene (Amp^R^) – generated as PCR amplicons of the Amp^R^ gene flanked by homology arms 50 to 600 bp in length – between the Kan^R^ and GFP genes on an established O-replicon (Supplementary Fig. 3). This process was highly efficient, yielding >10^8^ colonies per electroporation without any observed false-positive colonies (Supplementary Fig. 3). Amplicons with longer homology arms yielded more colonies per electroporation – 600 bp homologies led to a 3-fold increase in colony numbers relative to an amplicon bearing 50 bp homologies.

We also demonstrated the complete replacement of the sequence between the ITRs in an O-replicon using the λ Red recombination system with 600 bp homology arms (Supplementary Fig. 4a); albeit with a lower efficiency than insertions. Using this approach, we generated longer replicons (10-20 kb) with an efficiency that was two orders of magnitude higher than the efficiency for the *de novo* establishment of the same replicons by the direct electroporation of PCR amplicons; the two approaches require similar experimental effort (Supplementary Fig. 4a). Additionally, the replacement process was high fidelity, yielding few false-positive colonies (Supplementary Fig. 4b).

### Transformation of extracted O-replicons reveals the minimal synthetic replication operon

Next, we demonstrated that we could extract intact O-replicons from *E. coli*, by plasmid extraction protocols, and transform them into fresh cells harboring variants of the synthetic replication operon (Supplementary Fig. 5a). These experiments revealed that the expression of TP and O-DNAP are necessary and sufficient for establishing and maintaining the extracted O-replicon and that extracted replicons could be transformed with high efficiencies (>10^7^ colonies per µg of extracted replicon; Supplementary Fig. 5b). We suggest that the SSB is required to establish the naked O-replicons generated by PCR because it protects the freshly transformed replicon DNA from degradation. However, when the O-replicon is protected by TP, as it is when replicons are extracted from cells, SSB is no longer required for establishing the replicon in fresh cells. These observations suggested that once the O-replicon is established only TP and O-DNAP are required for its maintenance. Our experiments demonstrated that the three-component system, composed of TP, O-DNAP, and the O-replicon, constitutes the minimal orthogonal replication system for EcORep.

We also measured the growth of wild-type *E. coli* DH10B in comparison to strains harboring synthetic replication operons, with or without the SSB and DSB, supporting the replication of a Kan^R^-Cm^R^(Q38TAG) O-replicon. This revealed a negligible fitness burden for EcORep setups (Supplementary Fig. 6). By contrast, inducing the over-expression of the SSB and DSB or inducing increased replicon copy numbers led to increased doubling times and reduced maximum cell densities (Supplementary Fig. 7).

### O-replicon with large payloads via replicon-REXER

Generating O-replicons with large DNA payloads (>20 kb) would enable the evolution and biosynthetic pathways and other multigene systems. However, the generation of replicons with these payloads by transformation of amplicons or λ Red recombination on existing O-replicons in *E. coli*, will be exceptionally challenging or impossible. The transformation efficiency of single- or double-stranded DNA into *E. coli* decreases with the length of the DNA and this limits the observed efficiency of λ Red recombination and O-replicon establishment ^22^.

We sought to adapt REXER (Replicon EXcision Enhanced Recombination), which has been used to replace large sections of the genome with synthetic DNA, to introduce large DNA payloads into O-replicons ^22^. REXER relies upon the transformation of a bacterial artificial chromosome (BAC) harboring a sequence of interest flanked by homology arms, marker cassettes, and Cas9 cleavage sites ^23^. Transformation of the BAC and selection for its presence ensures that every cell in the experiment harbors a copy of the sequence to be recombined and thereby uncouples the efficiency of DNA delivery from the efficiency of recombination. The sequence of interest is then excised from the BAC using CRISPR/Cas9 and recombined into its target sequence, as guided by the homology arms, using the λ Red recombination machinery ^24^.

We designed a BAC for O-replicon-targeting REXER (replicon-REXER) consisting of 75 kb of human CFTR sequence with two flanking and one internal positive selection markers (+2, *kanR*; +3, *gmR*; +4, ampR), flanking homology arms 90 and 300 bp in length, and a dual positive-negative selection cassette in the BAC backbone (+1, *zeoR*; −1, *rpsL*) (Supplementary Fig. 8a). The target O-replicon was designed to harbor the necessary homology regions and an additional marker (+5, *specR*) for maintenance of the replicon. Importantly, the second homology region (HR2, 300 bp) was designed to generate an intact chloramphenicol-resistance gene (+6) only upon successful recombination (Supplementary Fig. 8a).

To perform the replicon-REXER, cells harboring the target O-replicon were transformed with the BAC carrying the sequence to be introduced onto the O-replicon, a plasmid encoding for the λ Red recombination machinery, and Cas9 was then transformed to initiate the excision of the sequence of interest from the BAC and its integration into the O-replicon. Selecting for the necessary positive markers (+2 to +6) and counter-selecting against the negative marker on the BAC backbone (−1) enabled the identification of cells harboring an intact O-replicon 77 kb in length. We passaged cells harboring this O-replicon and validated the stability of the replicon via next-generation sequencing. This revealed an even read depth across the replicon, with the exception of a 1058 bp sequence with 76% GC content which proved recalcitrant to Illumina sequencing and PCR (Supplementary Fig. 8b).

These experiments revealed that the PRD1 replication machinery used for EcORep can replicate sequences much longer than the PRD1 genome (15 kb). This suggested that PRD1genome size is not constrained by the replicative capacity of the PRD1 TP/DNAP system and might be constrained by other factors; for example, the space available in the phage capsid. We concluded that replicon-REXER is a method capable of establishing O-replicons that are longer than any sequence that has been replicated by any other orthogonal replication system to date.

### Directed evolution of error-prone O-DNAPs with less biased mutational spectra

We previously generated mutagenic O-DNAPs via rational design. These O-DNAPs exhibited moderate O-replicon mutation rates between 2.3 x 10^-8^ to 7.6 x 10^-6^ substitutions per base per generation (*μ*, s.p.b.) as measured via fluctuation tests ^25,26^. Here, we used directed evolution to generate highly mutagenic O-DNAPs with minimally biased mutational spectra. We generated multiple Cm^R^-Kan^R^ reporter O-replicons to select the desired O-DNAPs. In these reporters the Cm^R^ gene was inactivated via the introduction of one or two TAG stop codons or via mutation of the essential His193 codon CAT to the Asp codon GAT (Fig. 1). Reversion of the stop codon(s) to a sense codon or Asp193 to His via mutations introduced by a mutagenic O-DNAP resulted in a functional Cm^R^ gene that confers chloramphenicol resistance to the cell. This enabled selection of mutant O-DNAPs by selection on chloramphenicol.

**Fig. 1.**
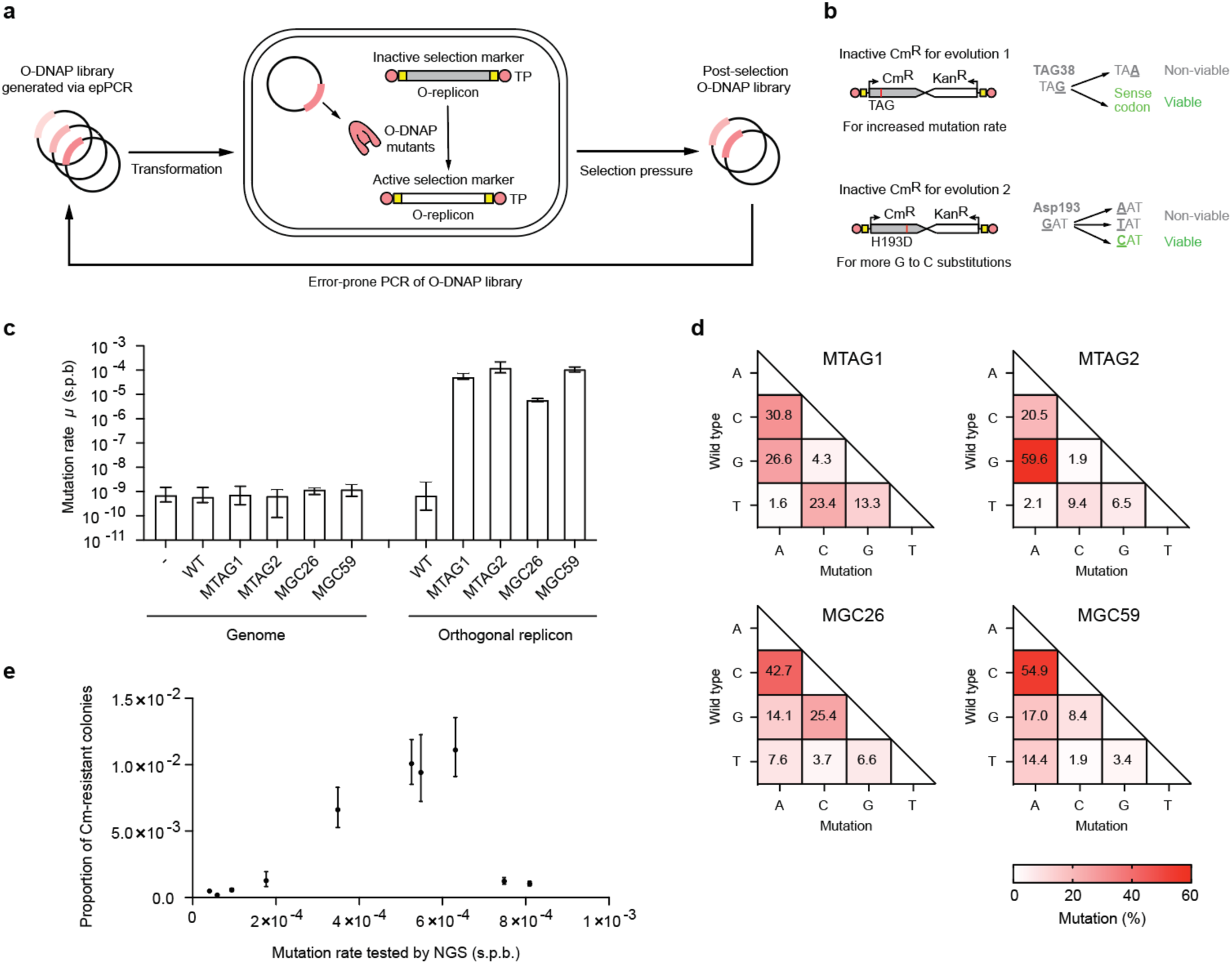
Directed evolution of error-prone O-DNAPs with less biased mutational spectra. **a**, Overview of the directed evolution of mutagenic O-DNAPs. **b**, Design of selection pressure for screening mutagenic O-DNAP variants. Multiple Cm^R^-Kan^R^ reporter O-replicons where the CmR gene was inactivated via the introduction of one or two TAG stop codons or via mutation of the essential His193 codon CAT to the Asp codon GAT were designed. Reversion of the stop codon(s) to a sense codon or Asp193 to His via mutations introduced by a mutagenic O-DNAP culminate in a functional CmR gene that confers chloramphenicol resistance to the cell. **c**, Determination of genomic or orthogonal replicon mutation rate (*μ*, s.p.b.) for the wild-type O-DNAP and the error-prone variants. The mutation rate was measured after 10 generations via fluctuation tests. For assessment of the orthogonal replicon mutation rate, we used an orthogonal replicon-encoded *Cm^R^* gene with a TAG stop codon at position 38 and the O-DNAP variant was expressed from a p15A plasmid via rhamnose induction (5 mM). For assessment of the genome mutation rate, we used a genomically-encoded *Cm^R^*gene with a TAG stop codon at position 38 and the O-DNAP variant was expressed from a p15A plasmid via rhamnose induction (5 mM). For all experiments n = 12, data are shown as median ± upper/lower 95% bounds. Supplementary Figs. 11 and 14 show the replicon mutation rates for other the error-prone O-DNAP variants. **d**, Mutational spectra for the four O-DNAP variants. n = 3, data are shown as mean. **e**, The critical error threshold for the genes under selection on the replicon (Cm and Kan, 2.1 kb total). n = 12, data are shown as mean ± s.d. Data from c and Supplementary Figs. 14 used to calculate the mutation rate were replotted in e.

Initially, we generated site-saturation mutagenesis libraries targeting residues in the exonuclease and DNAP domains and at the interface between the two, predicted from a combination of structural analysis and predicted homology to residues which had previously been reported to yield error-prone variants of the Φ29 DNAP and OrthoRep O-DNAP. However, this approach yielded variants with only a marginal increase in mutation rate (Supplementary Fig. 9). Thus, we proceeded with random, rather than targeted, mutagenesis of the O-DNAP for subsequent directed evolution campaigns.

We generated an O-DNAP library using error-prone PCR, with the rationally designed Y127A O-DNAP mutant as a template and transformed this library into cells harboring a TAG stop codon reporter O-replicon. We then passaged the resultant transformants for 10 generations to allow for the O-replicon mutagenesis to occur. Cells harboring mutagenic O-DNAP variants were then selected using chloramphenicol, the O-DNAP plasmid library was extracted from these cells, and the selected library was transformed into cells harboring the next reporter O-replicon for the next selection step (Supplementary Fig. 10). We performed three rounds consisting of two selection steps each and the output from each round was diversified via error-prone PCR prior to the next set of selections (Supplementary Fig. 10).

Ultimately, we identified an O-DNAP variant with just two mutations in addition to the parental Y127A mutation – Y347C and I367T (Fig. 1c, Supplementary Fig. 11a). Whereas the Y127A mutation is in the exonuclease domain, likely deactivating it, the other mutations are in the DNAP domain of the polymerase (Supplementary Fig. 11b). We named this O-DNAP mutant MTAG1 and found it to be highly error-prone and orthogonal, as assessed by fluctuation tests that rely on the reversion of a stop codon in a Cm^R^ gene on the O-replicon or on the genome (Fig. 1c). The mutation rate of MTAG1 is 5.11×10^-5^ s.p.b., corresponding to roughly one mutation per 10 generations (one passage) in a 2 kb gene. Another variant identified from these selections was MTAG2 (Y127F I317V K329R Y347C I367T P441Q), which had a mutation rate of 1.23×10^-4^ s.p.b. on the O-replicon without increasing genome mutagenesis (Fig. 1c). By contrast, the expression of an error-prone T7 DNAP variant resulted in a high level of genome mutagenesis (Supplementary Fig. 12) ^21^. By passaging cells harboring an O-replicon maintained by MTAG1 or MTAG2 for 20 generations, we determined the mutational spectra of the O-DNAPs using next-generation sequencing. We found that MTAG2 was highly biased for G:C to A:T mutations whereas MTAG1 had a less biased mutational spectrum (Fig. 1d).

For both O-DNAPs, MTAG1 or MTAG2, evolved via the TAG-inactivated Cm^R^ gene, we observed a low level of G:C to C:G transversion mutations. On this basis, we performed another directed evolution campaign using the H193D Cm^R^ gene (Fig. 1b). For this selection, only mutation of **G**AT to **C**AT results in cell survival. We generated an O-DNAP library using error-prone PCR, with the Y127A O-DNAP mutant as a template and transformed this library into cells harboring a H193D Cm^R^ reporter O-replicon. As before, we iterated rounds of selection and error-prone PCR (Fig. 1a; Supplementary Fig. 13).

Ultimately, we identified a series of mutagenic O-DNAPs with mutation rates between 1.53 x 10^-6^ to 1.23 x 10^-4^ s.p.b., as measured via fluctuation tests (Supplementary Fig. 14a). These identified O-DNAP variants had 6 to 13 mutations in addition to Y127A, distributed across both the exonuclease domain and the DNAP domain of the polymerase (Supplementary Fig. 14b). We further characterized two DNAPs, MGC26 (E97D, Y127A, G148S, L159I, F181L, A228V, N353K, I478T) and MGC59 (M81L, I102V, E121D, Y127A, G148S, L159I, N176K, F181L, H227Y, K297R, A419T, I478T); these DNAPs had mutation rates of 5.77 x 10^-6^ and 1.09 x 10^-4^ s.p.b., respectively, by fluctuation analysis, without mutating the genome (Fig. 1c). By passaging cells harboring an O-replicon maintained by MGC26 or MGC59 for 20 generations, we determined the mutational spectra of MGC26 and MGC59 using next-generation sequencing. We found that these O-DNAPs had mutational spectra with reduced bias (Fig.1d).

At high mutation rates fluctuation analysis may become a less accurate measurement of mutation rate, as mutations at sites in addition to the target codon site accumulate and contribute to the phenotype that is scored. We therefore further characterized the mutation rate of selected O-DNAPs on the Kan^R^-Cm^R^(Q38TAG) O-replicon by next-generation sequencing after 20 generations. O-DNAP variants MTAG1, MTAG2, MGC26, and MGC59 had mutation rates of 5.10 x 10^-5^, 7.49 x 10^-5^, 9.40 x 10^-5^, and 5.48 x 10^-4^ s.p.b. respectively by next-generation sequencing. These analyses demonstrated that the mutation rates of selected O-DNAPs were as high as 8.09 x 10^-4^ s.p.b (Fig. 1e, Supplementary Fig. 14c). Interestingly, we find that as the mutation rates of O-DNAPs rise from 5.91 x 10^-5^ s.p.b. to 6.31 x 10^-4^ s.p.b., as measured by next-generation sequencing, the fraction of Cm resistant colonies from Kan^R^-Cm^R^(Q38TAG) O-replicon increases, under selection for resistance to chloramphenicol and kanamycin. However, as the mutation rate increases further from 6.31 x 10^-4^ s.p.b. to 8.09 x 10^-4^ s.p.b. the number of resistant cells decreases. This observation is consistent with reaching the critical error threshold for the genes under selection on the replicon (Cm and Kan, 2.1 kb total) and suggests that the mutant O-DNAPs we have selected may span the critical error threshold for these genes (Fig. 1e).

Overall, our directed evolution experiments yielded O-DNAP variants with mutation rates that are at least two orders of magnitude higher than our previous variants. Moreover, some of these variants, such as MTAG1 and MGC26, have best-in-class mutational spectra among highly mutagenic DNAPs (∼10^-4^ s.p.b.) used in protein-primed orthogonal replication systems (Supplementary Fig. 15) - this will enable the less biased exploration of evolutionary pathways. The O-DNAPs we have discovered span the critical error threshold for the genes under selection. These observations suggest that we have discovered polymerases (with mutation rates just below the critical error threshold) that can mutate these genes as fast as possible without leading to catastrophic loss of gene function.

### System for the control of O-replicon mutagenesis

To control the mutagenesis during continuous evolution experiments, we used a strain where we had genomically integrated a synthetic replication operon with a WT O-DNAP under the control of an IPTG-inducible Ptac promoter. Additionally, we genomically integrated lacI, which represses the IPTG-inducible promoter, under the control of a constitutive promoter, ensuring that the basal O-replicon copy number remained low (approx. 3 to 5) in the absence of IPTG (lacI6; Supplementary Figs. 16 and 17). To induce O-replicon mutagenesis, we expressed TP and a mutagenic O-DNAP variant from a plasmid under the control of a stringent salicylic acid inducible promoter (Supplementary Fig. 18). This setup thus allows for straightforward control of mutagenesis, overcoming the need for additional dCas9-based repression of copy number which we utilized in the first iteration of EcORep ^14^.

### Accelerated continuous evolution of an ethanol assimilation pathway in *E. coli*

Next, we investigated whether we could use the improved, highly mutagenic EcORep system to continuously evolve multiple genes in a pathway for a new phenotype. We chose the *E. coli* ethanol assimilation pathway, consisting of an alcohol dehydrogenase (AdhP), an aldehyde dehydrogenase (MhpF), and a bifunctional aldehyde/alcohol dehydrogenase (AdhE) (Fig. 2a). To initiate these evolution experiments, we generated an O-replicon harboring the *adhE-adhP-mhpF* operon under the control of a constitutive PM4 promoter and passaged cells harboring this replicon in M9 minimal medium supplemented with 10% LB, 10 µM salicylic acid (to induce O-replicon mutagenesis mediated by the MTAG1 O-DNAP), 20 g L^−1^ ethanol, and decreasing concentrations of glucose (Supplementary Figs. 19 and 20). After five passages, we obtained pools of cells that grew to much higher maximum cell densities relative to the input control (Fig. 2b, Supplementary Fig. 21). We cloned different hits from three independent evolution replicates onto a standard circular plasmid (pSC101) and found that these clonal sequences facilitated strongly increased growth in the presence of 20 g L^−1^ ethanol (Fig. 2c, Supplementary Figs. 22-25).

**Fig. 2.**
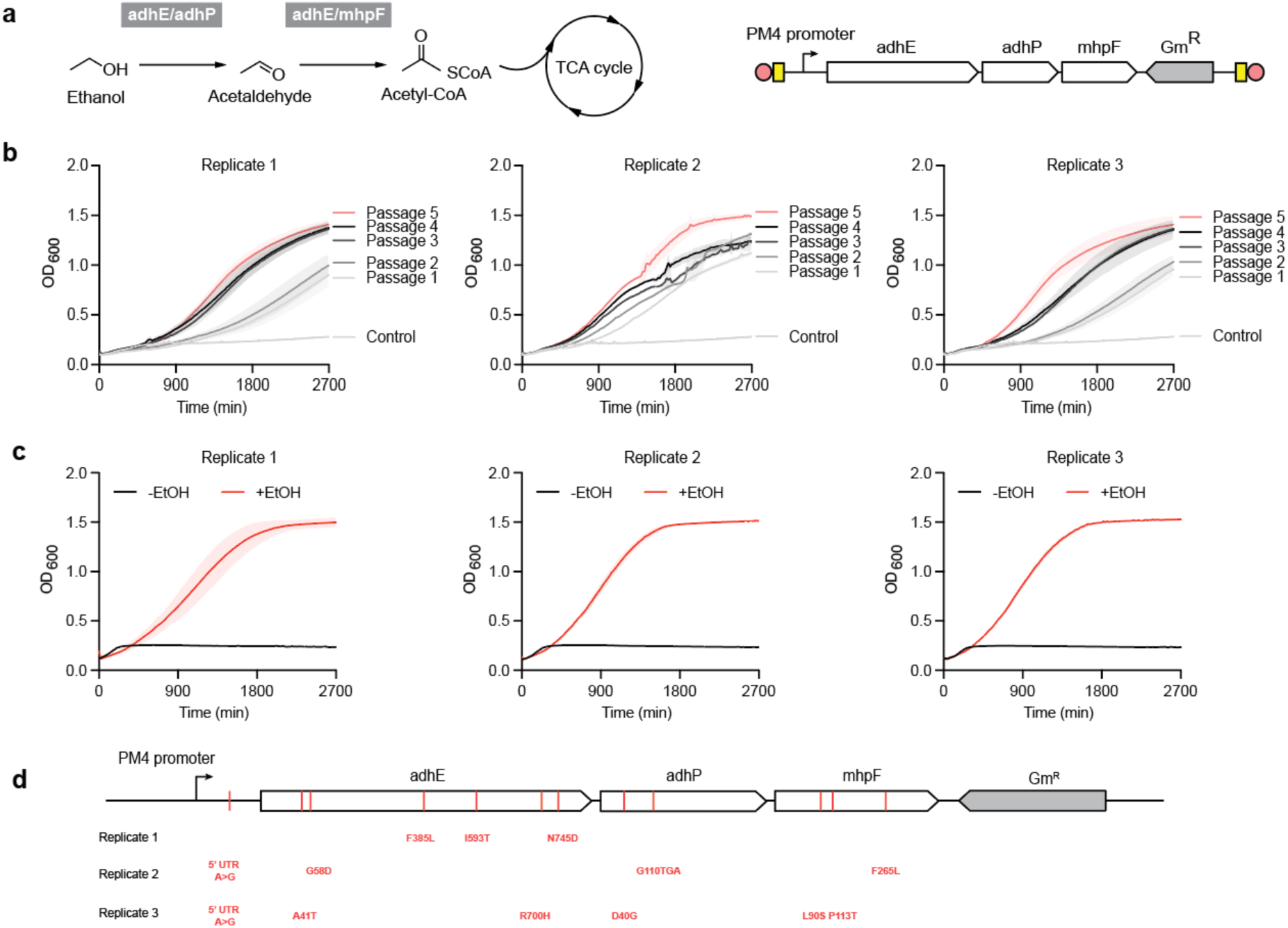
Accelerated continuous evolution of an ethanol assimilation pathway in *E. coli*. **a**, Design of an O-replicon encoding an ethanol assimilation pathway comprising genes encoding for three enzymes: aldehyde/alcohol dehydrogenase (AdhE), alcohol dehydrogenase (AdhP), and aldehyde dehydrogenase (MhpF). **b**, Growth curves of three evolved pools carrying the pathway-encoded orthogonal replicon after the indicated number of passages (n = 4, data are shown as mean ± s.d.). The same control was used for all three plots. The media were supplemented with 20 g L^-1^ ethanol. **c**, Validation of evolved pathways on a pSC101 plasmid. Shown is one example from each replicate; Supplementary Figs. 23-25 show other mutants (n = 4, data are shown as mean± s.d.). **d**, Non-synonymous and 5’-UTR mutations enriched in each replicate.

We found that mutations in the pathway were enriched within replicates but that there was minimal convergence of mutations between replicates; this suggested that functionally similar solutions were rapidly accessed through diverse genotypes. Over the three replicates, non-synonymous mutations were enriched across all three genes and the 5’-UTR (Fig. 2d, Supplementary Fig. 26). Notably, given the redundancy provided by AdhE, which acts as both an alcohol and aldehyde dehydrogenase, we found that in replicate two a stop codon had evolved in *adhP*.

### Establishing PRD1-based orthogonal replication in the fastest growing organism

The rate of continuous evolution via orthogonal replication depends on both the O-DNAP mutation rate and the generation time (i.e. doubling time). Therefore, we hypothesized that the fastest possible accelerated continuous evolution system for genes could be created by establishing an O-DNAP with a mutation rate of approximately 10^-4^ s.p.b for orthogonal replication in the organism with the most rapid generation time (Fig. 3a). *Vibrio natriegens* is the ideal candidate for this purpose as it has the fastest growth rate of any known organism ^27^ and is broadly compatible with commonly used molecular biology tools. We therefore set out to establish the PRD1-based orthogonal replication system in *V. natriegens*.

**Fig. 3.**
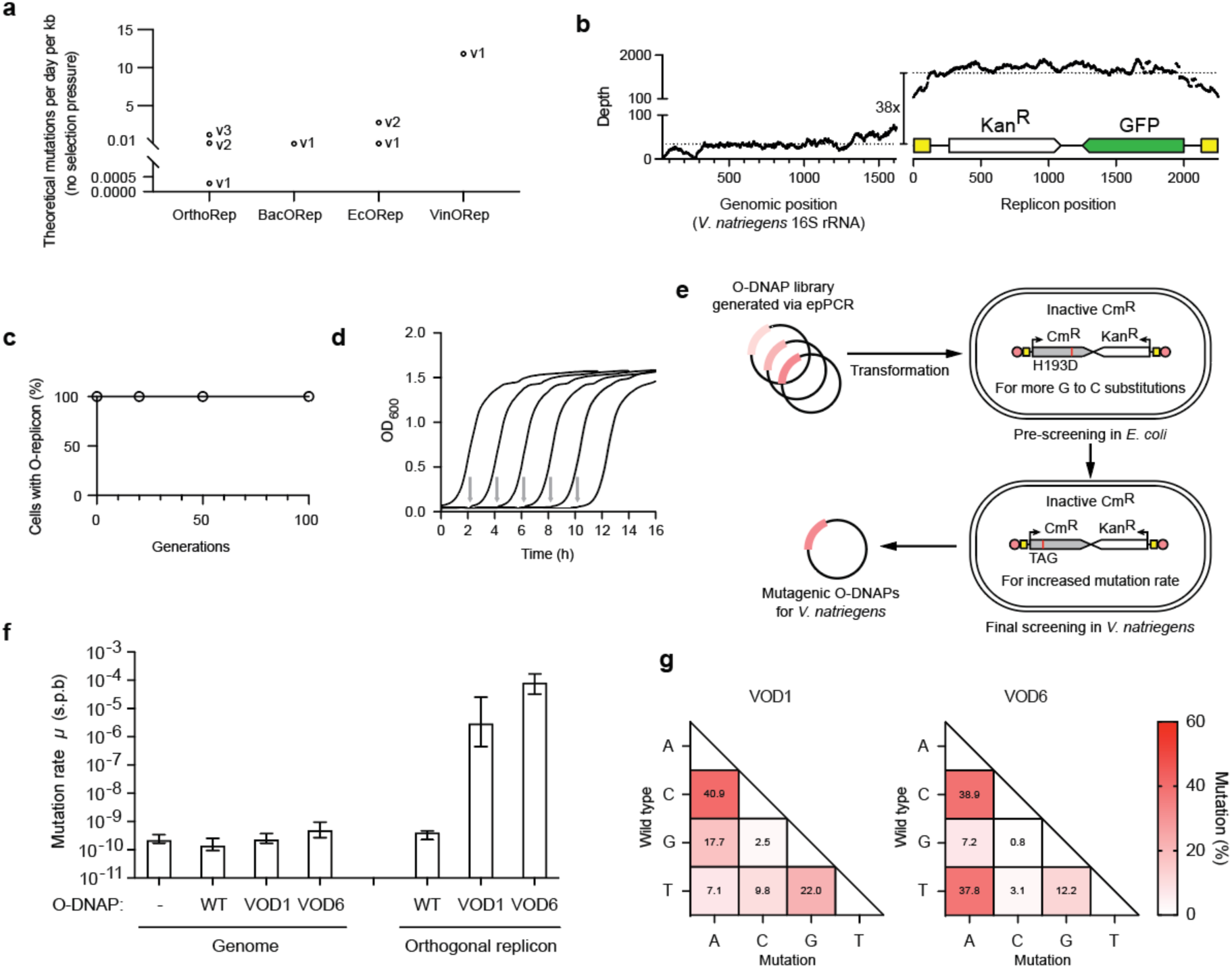
Establishing a synthetic orthogonal DNA replication system in *Vibrio natriegens*. **a**, Summary of theoretical mutations per day per kilobase in the absence of selection pressure across all orthogonal replication systems. The mutation rates and doubling times used in the calculations were obtained from the literature (Extended Data Table 1). **b**, The synthetic replication operon was under the control of an IPTG inducible promoter (PtacIPTG). The Kan^R^-GFP replicon consists of flanking 110 bp ITR, a kanamycin resistance gene, and a GFP gene. Shown is Illumina sequencing read coverage. **c**, Orthogonal replicons are stably maintained in *V. natriegens* (n = 3, data are shown as mean ± s.d.). **d**, Growth curves of wild-type *V. natriegens* and *V. natriegens* harboring a replicon (n = 8, data are shown as mean ± s.d.). **e**, Overview of the directed evolution of mutagenic O-DNAP for *V. natriegens*. **f**, Determination of genomic or orthogonal replicon mutation rate (*μ*, s.p.b.) for the wild-type O-DNAP and the evolved variants in *V. natriegens*. The mutation rates were measured after 10 generations via fluctuation tests. For assessment of the orthogonal replicon mutation rate, we used an orthogonal replicon-encoded *Cm^R^* gene with a TAG stop codon at position 38 and the O-DNAP variant was expressed from a p15A plasmid under the control of a constitutive promoter. For assessment of the genome mutation rate, we used the host *rpoB* gene as a marker to detect mutations that result in rifampicin resistance and the O-DNAP variant was expressed from a p15A plasmid under the control of a constitutive promoter. For all experiments n = 12, data are shown as median ± upper/lower 95% bounds. Supplementary Fig. 33 shows the replicon mutation rates for other the error-prone O-DNAP variants. **g**, Mutational spectra for the O-DNAP variants.

To express the minimal synthetic replication operon (consisting of the TP and O-DNAP) in *V. natriegens,* we used a plasmid with an origin of replication that has a broad host range (RSF1010). We then transformed replicons extracted from *E. coli* into *V. natriegens* (Vmax X2) harboring the operon-encoding plasmid (Supplementary Fig. 27a). We confirmed the establishment of the O-replicon in *V. natriegens* via genotyping and next-generation sequencing (Fig. 3b; Supplementary Fig. 27b). We also passaged cells transformed with the O-replicon for 100 generations and found that the O-replicon was stably maintained (Fig. 3c; Supplementary Fig. 28). Furthermore, we measured the growth of the strain with or without the synthetic replication operon and an O-replicon and found that these components imposed only a minor fitness burden on the cells (Supplementary Fig. 29). We were able to reproducibly passage (100-fold dilutions) *V. natriegens* harboring the O-replicon five times in a 12-hour timeframe, totaling 40 generations (Fig. 3d).

In additional experiments, we also established the PRD1-based replication system in the industrially-relevant *Pseudomonas putida* strain KT2440 (Supplementary Figs. 30-32). Thus, the PRD1-based orthogonal replication system can be established, beyond *E. coli*, in other Gram-negative species of biotechnological significance.

We initially tested the performance of O-DNAP variants MTAG2 and MGC59 which were highly mutagenic in *E. coli*. However, while these O-DNAPs were mutagenic in *V. natriegens*, with a maximum mutation rate on the replicon of 3.21 x 10^-6^ s.b.p. (by fluctuation analysis, Supplementary Fig. 33), they were not as mutagenic as in *E. coli*. This suggested that unknown host specific effects (e.g., endogenous DNA repair activity, O-DNAP expression level, endogenous nucleotide pool levels) may lead to differences in observed net mutation between distinct hosts.

We decided that the most parsimonious route to discovering a highly mutagenic O-DNAP in *V. natriegens* was to evolve new mutagenic O-DNAPs directly in *V. natriegens*. Following a pre-screening enrichment step in *E. coli*, where plasmid transformation efficiencies are higher, we selected O-DNAPs in *V. natriegens* using a TAG-inactivated Cm^R^ O-replicon reporter (Fig. 3e). Ultimately, we identified a series of mutagenic O-DNAPs. These O-DNAPs exhibited O-replicon mutation rates between 1.16 x 10^-6^ to 8.18 x 10^-5^ s.p.b. as measured via fluctuation tests in *V. natriegens* (Supplementary Fig. 33).

We further characterized two O-DNAPs: VOD1 (F84L, Y127A, G148S, L159I, I478T, K504I) and VOD6 (V9M, F84L, Y127A, G148S, L159I, I478T, D489E, K501I). The mutation rate of these O-DNAPs on the O-replicon were 2.91 x 10^-6^ and 8.18 x 10^-5^ s.p.b., respectively, by fluctuation analysis (9.84 x 10^-5^ and 9.36 x 10^-5^ s.p.b by next-generation sequencing); these O-DNAPs do measurably not mutate the *V. natriegens* genome (Fig. 3f). By passaging cells harboring an O-replicon maintained by VOD1 or VOD6 for 20 generations, we determined the mutational spectra of the O-DNAPs using next-generation sequencing (Fig. 3g).

Our observations confirmed that the PRD1-based replication system is orthogonal in *V. natriegens* and that we can establish versions of this system that mutate genes on the replicon at approximately 10^-4^ s.p.b with a well distributed spectrum of mutations. These data suggested that we had created the fastest possible continuous evolution system for gene length DNA in which mutagenic replication just below the critical error threshold of genes tested is realized in the fastest growing organism. We named the *V. natriegens* Orthogonal Replication system VinORep.

### Highly accelerated continuous evolution

Next, we demonstrated that VinORep enables highly accelerated continuous evolution. We transformed an O-replicon encoding for *tetA*, a tetracycline resistance-conferring gene, into *V. natriegens* cells harboring both the synthetic replication operon plasmid and a separate plasmid expressing the mutagenic VOD6 O-DNAP. We then evolved *tetA* to confer tigecycline resistance in a single day of passaging (Fig. 4a). We performed four passages in increasing concentrations of tigecycline in 16 hours, allowing the final passage to grow overnight; each sequence of passages was performed in four replicates. The following day, we cloned the evolved sequences from each of the four replicates into a circular plasmid backbone in *E. coli*. The pooled plasmids were then transformed back into *V. natriegens* to ensure that each cell contained a single clonal sequence and that the plasmid copy number was consistent across all mutants. We sequenced 20 colonies from each of the four evolution replicates, and this revealed a high number of mutations on the replicons (mean= 10.82 mutations/replicon). The most mutated clone accumulated 30 mutations across its 2.2 kb sequence (Fig. 4b).

**Fig. 4.**
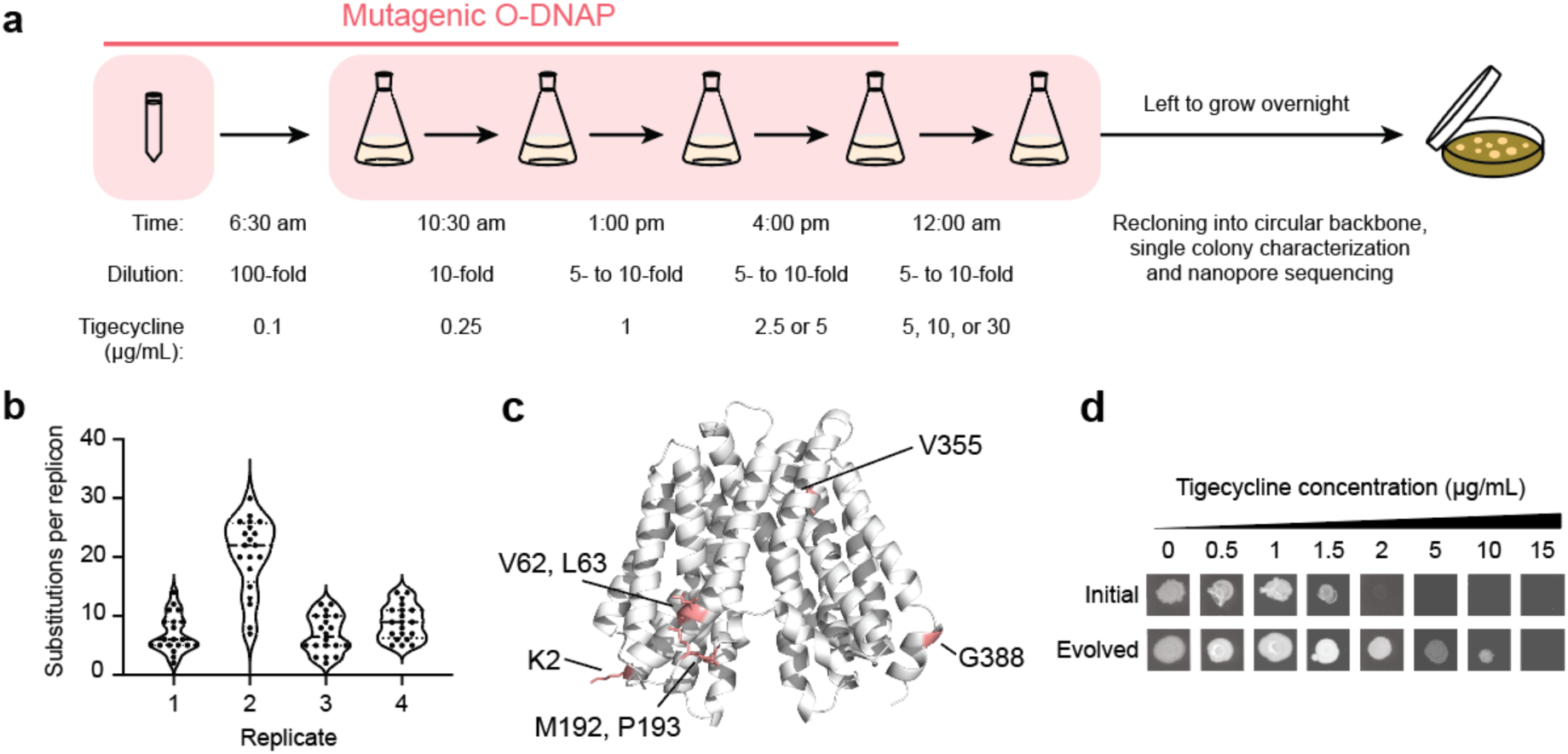
Highly accelerated continuous evolution in *Vibrio natriegens*. **a**, Overview of the *tetA* accelerated continuous evolution. **b**, Substitutions per replicon after evolution. n = 20, data are shown as violin plots with interquartile range. **c**, AlphaFold2 model of TetA. The residues that had mutated in at least two independent evolution replicates or have been reported in the previous *tetA* evolution experiments in *E. coli* are shown in stick representation. **d**, Validation of evolved *tetA* on a p15A plasmid. Shown is Rep3_2 from the replicate 3 pool; Supplementary Fig. 34 shows the activity of other mutants.

Sequencing evolved clones identified mutations in the promoter and 5′-untranslated region (5′-UTR), as well as a number of synonymous and nonsynonymous mutations in the open reading frame. We observed a 5′-UTR mutation, G128T, across all replicates, which increased the RBS calculator predicted translation rate of *tetA* (from 792.8 a.u. to 2045.7 a.u.) ^28^. We also identified several mutations that have been reported in the previous *tetA* evolution experiments in *E. coli*, including M192I (replicate 1), P193H (replicate 3), V355F (replicates 1 and 3), and a range of C-terminal mutations (A384 to T396), including the previously reported G388C mutation, were found in replicates 1, 2, and 4. In addition to previously described mutations, we identified a new N-terminal coding sequence mutation, K2N or K2I, in both replicates 1 and 2. Furthermore, motif substitutions from “APVLG” to “APVIG” or “APFLG” were detected within the replicates; both variants have been observed in naturally occurring *tetA* mutants (Fig. 4c).

The evolved *tetA* genes conferred tigecycline resistance to 10 μg/ml, whereas the parent *tetA* gene conferred resistance to 1.5 μg/ml (Fig. 4d, Supplementary Fig. 34). Thus, VinORep enables the selection of new phenotypes within a day and realizes the advantages of combining the mutation rates approaching the limit of molecular evolution for genes with the fastest growing organism to realize the biological speed limit for continuous gene scale evolution using orthogonal replication systems (Fig. 3a).

## Discussion

We have upgraded key aspects of EcORep - an orthogonal DNA replication system that operates in the workhorse of biotechnology, *E. coli*. We developed and optimized methodologies that enable the efficient establishment, engineering, and transformation of O-replicons, streamlining workflows for setting up new accelerated evolution experiments. In doing so, we discovered that just two genes, encoding for the O-DNAP and TP, are needed to stably maintain the O-replicon. Using replicon-REXER, we established a 77 kb O-replicon which constitutes the longest orthogonally replicating episome to date and suggests that EcORep may enable the evolution of large gene clusters or entire genomic sections.

We utilized directed evolution to generate highly mutagenic O-DNAP variants which are less mutationally biased than those of previous systems and mutate the O-replicon about a million fold faster than the genomic mutation rate. Importantly, these O-DNAPs exhibited a range of mutation rates and the most mutagenic variants led to deleterious levels of gene hypermutation as determined by elevated loss of gene function. This experimental observation of error catastrophe at the single-gene level suggests that the evolved O-DNAP mutation rates are distributed around the critical error threshold for the genes tested. We utilize the highly mutagenic and mutationally less biased EcORep system to rapidly evolve an ethanol assimilation pathway for improved performance.

Leveraging the methodological developments in *E. coli*, the components that drive EcORep were used to drive O-replicon establishment and maintenance in other Gram-negative bacteria. VinORep, which operates in the fastest growing organism *V. natriegens*, combines the high number of mutations per generation of evolved O-DNAPs with rapid generation time, enabling an order of magnitude increase in the accumulation of mutations relative to the *E. coli* system. This system enables the in vivo continuous evolution of genes at the biological speed limit. We demonstrate that this enables highly accelerated continuous evolution by rapidly evolving *tetA* to confer tigecycline resistance in a single day of passaging.

The systems we have developed will provide less biased access to diverse sequence space at an unprecedented pace, enabling the rapid exploration of fitness landscapes, dissection of evolutionary pathways, and pre-emptive discovery of resistance mutations. We expect that these approaches will enable the accelerated evolution of diverse genes, multi-gene pathways, and sections of genomes for new and improved functions.

## Methods

### Strains

All *E. coli* strains used in this study are derived from *E. coli* DH10B. The strain with a genomically integrated synthetic replication operon controlled by an IPTG inducible promoter (PtacIPTG) was described previously ^14^. We integrated a *lacI* gene under the control of different promoters into the genome of *E. coli* through λ Red knock-in, as described before (pEcCas9). The concentrations of the corresponding antibiotic used in this study (unless otherwise specified) include: kanamycin resistance gene (*Kan^R^*, 50 µg mL^-1^), chloramphenicol resistance gene (*Cm^R^*, 20 µg mL^-1^), tetracycline resistance gene (*tetA*, 10 µg mL^-1^), apramycin resistance gene (*Apm^R^*, 50 µg mL^-1^), zeocin resistance gene (*Zeo^R^*, 25 µg mL^-1^), spectinomycin resistance gene (*Spec^R^*, 50 µg mL^-1^), hygromycin resistance gene (*Hyg^R^*, 200 µg mL^-1^) and ampicillin resistance gene (*Amp^R^,* 50 µg mL^-1^).

The *Vibrio natriegens* strain used was Vmax X2. The *Pseudomonas putida* strain used was KT2440.

### Construction of orthogonal replicons

Linear replicons used for electroporation were obtained by overlap extension PCR (PrimeSTAR Max DNA Polymerase, TAKARA, Tokyo, Japan). Specifically, four or more DNA fragments with overlaps, including left replication origin, antibiotic resistant genes, genes of interest and right replication origin were first obtained by PCR and gel purified. Then a single primer (without modification) was used to amplify the whole length orthogonal replicon by binding to both inverted repeat ends of the template.

Primer sequence used:

5’-ggggatacgtgcccctccccacctacccgcgcccctaacatttttatttccgtc-3’

### Construction of circular plasmids

To perform λ Red recombination experiments, we designed a circular plasmid pEcCas9_pH based on the pSC10 replication origin, a hygromycin resistance gene (*Hyg^R^*), a negative selection marker (*pheS*\*), the λ Red components under the control of an arabinose-inducible promoter and a CRISPR-Cas9 gene under the control of a constitutive promoter. Plasmids overexpressing an error-prone O-DNAP or a combination of both TP and error-prone O-DNAP under the control of a rhamnose inducible promoter were designed based on a p15A replication origin and an apramycin resistant gene (*Apm^R^*).

All circular plasmids were constructed using HiFi Gibson Assembly (New England Biolabs, Massachusetts, United States) from multiple fragments.

### Electroporation protocol for establishing O-replicons

First, colonies harboring the helper plasmid pFR160GB and a genomically-integrated synthetic replication operon were inoculated into 5 ml 2XTY media in a 50 mL falcon tube and grown at 37 °C while shaking (220 rpm) for 12 h. Second, 2 mL of the resulting culture was used to inoculate a 2 L shake flask containing 200 mL 2XTY media and grown at 37 °C while shaking (220 rpm) to and OD_600_ of ∼0.3. Arabinose was then added to the culture to a final concentration of 10 mM. Cells were then induced for different times (15 min, 30 min or 60 min) and then chilled on ice while shaking (100 rpm) for 10 min and then spun down (4500 xg, 3 min, 4 °C) and washed in 50 mL pre-chilled 10% (v/v) glycerol twice. After washing, cell pellets were finally resuspended in 1 mL of ice-cold 10% (v/v) glycerol and aliquots of 100 μL were used for orthogonal replicon electroporation. For electroporation, 5 μL purified orthogonal replicon PCR product (around 500 ng μL^−1^, in water) was mixed with 50 μL competent cells. The mixture was added to an ice-cold electroporation cuvette (2 mm gap; SLS scientific), and incubated on ice for 15 min. The electroporation was performed using Eppendorf e-porator (2500 V) and 1 mL of pre-warmed (37 °C) SOB media was then immediately added to the cuvette. After incubating the mixture at 37 °C while shaking (220 rpm) for 2 h, all cells were plated on an agar plate with the appropriate antibiotics.

### Engineering O-replicons *in vivo* using the λ Red recombination system

For the insertion of an ampicillin resistance gene, the PCR amplicons of the *Amp^R^* gene flanked by homology arms 50 and 100 bp in length were generated directly by primers binding to both ends of the *Amp^R^* gene. The PCR amplicons of the *Amp^R^* gene flanked by homology arms 200 to 600 bp were obtained by overlap extension PCR (Phanta Flash Master Mix, Vazyme, Nanjing, China). Specifically, three DNA fragments with overlaps, including a left homology arm, Amp^R^ gene, and a right homology arm, were first obtained by PCR and gel purified. Then two primers were used to amplify the whole length by binding to both ends of the template.

All other PCR products used for engineering O-replicons by λ Red recombination also were obtained by overlap extension PCR.

For the transformation, first, colonies harboring the plasmid pEcCas9_pH encoding λ Red components under the control of an arabinose-inducible promoter and a O-replicon were inoculated into 5 mL 2XTY media in a 50 mL falcon tube and grown at 37 °C while shaking (220 rpm) for 12 h. Second, 2 mL of the resulting culture was used to inoculate a 2 L shake flask containing 200 mL 2XTY media and grown at 37 °C while shaking (220 rpm) to and OD_600_ of ∼0.3. Arabinose was then added to the culture to a final concentration of 10 mM. Cells were induced for 30 min and then chilled on ice while shaking (100 rpm) for 10 min and then spun down (4500 xg, 3 min, 4 °C) and washed in 50 mL pre-chilled 10% (v/v) glycerol twice. After washing, cell pellets were finally resuspended in 1 mL of ice-cold 10% (v/v) glycerol and aliquots of 100 μL were used for orthogonal replicon electroporation. For electroporation, 5 μL purified orthogonal replicon PCR product (around 100 ng μL^−1^, in water) was mixed with 50 μL competent cells. The mixture was added to an ice-cold electroporation cuvette (2 mm gap; SLS scientific), and incubated on ice for 15 min. The electroporation was performed using Eppendorf e-porator (2500 V) and 1 mL of pre-warmed (37 °C) SOB media was then immediately added to the cuvette. After incubating the mixture at 37 °C while shaking (220 rpm) for 2 h, all cells were plated on an agar plate with appropriate antibiotics.

### Replicon-REXER

We first designed a bacterial artificial chromosome (BAC) consisting of 75 kb of human CFTR sequence with two flanking and one internal positive selection markers (+2, *kanR*; +3, *gmR*; +4, ampR), flanking homology arms 90 and 300 bp in length, and a dual positive-negative selection cassette in the BAC backbone (+1, *zeoR*; −1, *rpsL*). The BAC was obtained by engineering a previously reported 200 kb BAC ^23^ by λ Red recombination. Then we transformed this BAC into a strain harboring a Spec^R^-mKate2 O-replicon with corresponding homology arms.

To perform the replicon-targeting REXER (replicon-REXER) experiment, first, colonies harboring the BAC and the O-replicon were inoculated into 5 mL 2XTY media in a 50 mL falcon tube and grown at 37 °C while shaking (220 rpm) for 12 h. Second, 5 mL of the resulting culture was used to inoculate a 2 L shake flask containing 200 mL 2XTY media and grown at 37 °C while shaking (220 rpm) to and OD_600_ of ∼0.3. Arabinose was then added to the culture to a final concentration of 10 mM. Cells were induced for 60 min and then chilled on ice while shaking (100 rpm) for 10 min and then spun down (4500 xg, 3 min, 4 °C) and washed in 50 mL pre-chilled 10% (v/v) glycerol twice. After washing, cell pellets were finally resuspended in 1 mL of ice-cold 10% (v/v) glycerol and aliquots of 100 μL were used for orthogonal replicon electroporation. For electroporation, 5 μL purified plasmid pEcCas9_PH (around 200 ng μL^−1^, in water) was mixed with 50 μL competent cells. The mixture was added to an ice-cold electroporation cuvette (2 mm gap; SLS scientific), and incubated on ice for 15 min. The electroporation was performed using Eppendorf e-porator (2500 V) and 1 mL of pre-warmed (37 °C) SOB media was then immediately added to the cuvette. After incubating the mixture at 37 °C while shaking (220 rpm) for 1 h, we added arabinose and hygromycin and incubated for another 3 h. All cells were diluted and plated on an agar plate with corresponding antibiotics.

### Orthogonal replicon extraction from cells

Extracted O-replicons could be transformed with a much higher efficiency than PCR products and without needing to provide a helper plasmid. To extract these replicons, 10 mL overnight culture of a strain harboring a synthetic replicon was pelleted by centrifugation (4500 x g, 10 min). Next, we used the QIAprep Spin Miniprep Kit to extract the replicon, following the manufacturer’s guidelines. We used Millipore-filtered water for the elution and heated the columns to which the water was added at to 50 °C for 10 min prior to the final centrifugation. All extracted replicons were stored at −20 °C.

### Electroporation protocol for transformation of extracted O-replicons into *E. coli*

First, colonies with a genomically-integrated synthetic replication operon were inoculated into 5 mL 2XTY media in a 50 mL falcon tube and grown at 37 °C while shaking (220 rpm) for 12 h. Second, 5 mL of the resulting culture was used to inoculate a 2 L shake flask containing 200 mL 2XTY media and grown at 37 °C while shaking (220 rpm) to and OD_600_ of ∼0.4. Cells were then chilled on ice while shaking (100 rpm) for 10 min and then spun down (4500 xg, 3 min, 4 °C) and washed in 50 mL pre-chilled 10% (v/v) glycerol twice. After washing, cell pellets were finally resuspended in 1 mL of ice-cold 10% (v/v) glycerol and aliquots of 100 μL were used for orthogonal replicon electroporation. For electroporation, 5 μL extracted O-replicons (around 100 ng μL^−1^, in water) was mixed with 50 μL competent cells. The mixture was added to an ice-cold electroporation cuvette (2 mm gap; SLS scientific), and incubated on ice for 15 min. The electroporation was performed using Eppendorf e-porator (2500 V) and 1 mL of pre-warmed (37 °C) SOB media was then immediately added to the cuvette. After incubating the mixture at 37 °C while shaking (220 rpm) for 2 h, all cells were plated on an agar plate with appropriate antibiotics.

### Electroporation protocol for transformation of extracted O-replicons to *V. natriegens*

First, colonies with a plasmid with a RSF1010 origin encoding the synthetic replication operon were inoculated into 5 mL 2XTYV (2XTY with V2 salts) media in a 50 mL falcon tube and grown at 37 °C while shaking (220 rpm) for 4 h. Second, 2 mL of the resulting culture was used to inoculate a 2 L shake flask containing 200 mL 2XTYV media with 0.1 mM IPTG and grown at 37 °C while shaking (220 rpm) to and OD_600_ of ∼0.6. Cells were then chilled on ice while shaking (100 rpm) for 10 min and then spun down (4500 xg, 3 min, 4 °C) and washed in 50 mL pre-chilled 1 M sorbitol four times. After washing, cell pellets were finally resuspended in 1 mL of ice-cold 10% (v/v) glycerol and aliquots of 100 μL were used for orthogonal replicon electroporation. For electroporation, 5 μL extracted O-replicons (around 200 ng μL^−1^, in water) was mixed with 50 μL competent cells. The mixture was added to an ice-cold electroporation cuvette (2 mm gap; SLS scientific), and incubated on ice for 15 min. The electroporation was performed using Eppendorf e-porator (2500 V) and 1 mL of pre-warmed (37 °C) 2XTYV media was then immediately added to the cuvette. After incubating the mixture at 37 °C while shaking (220 rpm) for 2 h, all cells were plated on an agar plate with appropriate antibiotics.

### Directed evolution of error-prone O-DNAPs

First, we generated multiple Kan^R^-Cm^R^ reporter O-replicons for two independent rounds of O-DNAP evolution (evolution_1 and evolution_2). In these reporters the Cm^R^ gene was inactivated via the introduction of one or two TAG stop codons for evolution_1 or via mutation of the essential His193 codon CAT to the Asp codon GAT for evolution_2. Reversion of the stop codon(s) to a sense codon or Asp193 to His via mutations introduced by a mutagenic O-DNAP culminate in a functional Cm^R^ gene that confers chloramphenicol resistance to the cell. This enabled selection of mutant O-DNAPs by selection on chloramphenicol.

For evolution_1, we first cloned the O-DNAP library by assembling an error-prone PCR amplified O-DNAP fragment generated using GeneMorph II (Agilent, Santa Clara, United States.), into a p15A plasmid encoding terminal protein under the control of a rhamnose promoter. The O-DNAP variants were cloned in the same gene cluster as the terminal protein. We then introduced a Kan^R^-Cm^R^(Q38TAG) orthogonal replicon into cells containing a genomically-encoded synthetic replication operon with a wild-type (WT) DNAP. We transformed these cells with the plasmid pRT19 encoding an arabinose inducible dCas9, which upon addition of arabinose represses the synthetic replication operon, and the O-DNAP library. All cells were inoculated into 250 mL 2XTY media with kanamycin, tetracycline, ampicillin, 10 mM arabinose and 10 mM rhamnose, and grown at 37 °C while shaking (220 rpm) for 12 h. Then 50 ml of the culture was inoculated into 250 mL 2XTY media with kanamycin, tetracycline, ampicillin and chloramphenicol, and grown at 37 °C to dense (∼OD_600_ = 5) while shaking (220 rpm) to enrich the target O-DNAP variants. The culture was diluted and plated on plates with or without 20 µg ml^-1^ chloramphenicol for sequencing verification. We used the QIAprep Spin Miniprep Kit to extract the O-DNAP library. Before transforming the extracted DNA into cells for further enrichment, we performed gel purification to remove O-replicons that were co-extracted by the kit. We then transformed the purified DNA into cells and did the same experiment for the mutagenic O-DNAP enrichment. After the first round of enrichment, the purified DNA was used as a template for the next rounds of error-prone PCR and selection as shown in Supplementary Fig. 10.

For evolution_2, we first cloned the O-DNAP library by assembling an error-prone PCR amplified O-DNAP fragment generated using GeneMorph II (Agilent, Santa Clara, United States.), into a p15A plasmid, placing the O-DNAP variants under the control of the rhamnose promoter. We then introduced a Kan^R^-Cm^R^(H193D) orthogonal replicon into cells containing a genomically-encoded synthetic replication operon with a wild-type (WT) DNAP. In this case, we did not overexpress the terminal protein or use the plasmid pRT19 encoding an arabinose-inducible dCas9, because we previously showed that overexpression of O-DNAP alone does not change copy number. We transformed these cells with the O-DNAP library. All cells were inoculated into 250 mL 2XTY media with kanamycin, ampicillin and 10 mM rhamnose, and grown at 37 °C while shaking (220 rpm) for 12 h. Then 50 mL of the culture was inoculated into 250 mL 2XTY media with kanamycin, ampicillin and chloramphenicol, and grown at 37 °C to dense (∼OD_600_ = 5) while shaking (220 rpm) to enrich the target O-DNAP variants. The culture was diluted and plated on plates with or without 20 µg ml^-1^ chloramphenicol for sequencing verification. And we used the QIAprep Spin Miniprep Kit to extract the O-DNAP library. Before transforming the extracted DNA into cells for further enrichment, we performed gel purification to remove O-replicons that were co-extracted by the kit. We then transformed the purified DNA into cells and did the same experiment for the mutagenic O-DNAP enrichment. After the first round of enrichment, the purified DNA was used as a template for the next rounds of error-prone PCR and selection as shown in Supplementary Fig. 13.

### Orthogonal replicon stability test in *V. natriegens* and *P. putida*

To test the stability of the orthogonal replicon in *V. natriegens* and *P. putida*, the cells were inoculated into a 24-well plate containing 3 mL 2XTY, with corresponding antibiotics, per well and grown at 37 °C with shaking (220 rpm) for 12 h. The culture, regarded as generation 0, was then inoculated into the next well containing 3 mL 2XTY, with the corresponding antibiotics, at a dilution factor of 1/1000 (10 generations) and cultured for 12 h (∼2^10^-fold increase in cells, generation 10). The inoculation and culturing steps were repeated 10 times, until generation 100. All the intermediate samples were streaked on agar plates, and 12 colonies were picked from each replicate for genotyping to verify the presence of the O-replicon.

### GFP expression measurements

To test the GFP expression levels of cells harboring orthogonal replicons, we first inoculated all strains into 96-well plates containing 200 μL 2XTY media with the corresponding antibiotics per well and grew them at 37 °C while shaking (750 rpm) for 12 h. Then we inoculated 10 μL of all the cultures into 190 μL 2XTY media with all corresponding antibiotics and the different concentrations of inducers. The plates were incubated at 37 °C while shaking (750 rpm) for 12 h, and we diluted 50 μL of the samples with100 μL of PBS in 96-well flat-bottom clear plates, and measured GFP fluorescence (λ_ex_: 485 nm; λ_em_: 520 nm) and OD_600_ using a PHERAstar FS plate reader (BMG Labtech, Ortenberg, Germany). In all cases, we report the GFP fluorescence normalized by OD_600_.

### Growth measurements

Bacterial colonies were grown overnight at 37 °C in 2xYT with the relevant antibiotics in a 96-well plate. Overnight cultures were diluted 1:100 and monitored for growth in a 200 µL volume in a clear flat bottom 96-well plate. Measurements of OD_600_ were taken every 5 min on a Agilent BioTek LogPhase 600 set at 37 °C with shaking. Doubling times were calculated by the AMIGA fitness analysis software ^29^.

### Plasmid copy number assay

For all *E. coli* samples, we first spun down and resuspended all cells in water and adjusted them to OD_600_ ∼1. Then a lysis buffer (QuickExtract DNA Extraction Solution) was used to lyse all cells, and we used the lysate as template for quantitative real-time PCR (qPCR) to test target DNA copy number. The reference gene was the *dxs* gene on the genome, as it has been shown to be a stable one-copy reference in *E. coli*. The target gene used was *kanR*. We used a fused DNA template containing the *dxs* gene and *kanR* gene to make sure that the copy number of the two genes was equal. We used qPCR to determine the DNA copy numbers for all samples with a Vii 7 Real-Time PCR System with 384-Well Block (Thermo Fisher, Massachusetts, United States).

Primer sequences used:

*dxs*: 5’-cttcatcaagcggtttcaca-3’ and 5’-cgagaaactggcgatcctta-3’

*kan^R^*: 5’-catggcaaaggtagcgttgcc-3’ and 5’-ccatgcatcatcaggagtacg-3’

For all *V. natriegens* and *P. putida* samples, copy number was determined using Illumina sequencing by comparing read-depth differences between the replicon and the genomic 16S rRNA gene.

### Mutation rate measurements

We used Luria-Delbrück fluctuation analysis to measure the genomic mutation rate of all strains ^26^. To measure the *E. coli* genomic mutation rate, first, we introduced a Cm^R^(Q38TAG) gene that contains an amber stop codon (TAG) at position 38 into the genome of *E. coli* DH10B. The insertion site was adjacent to the *lacI* gene, distal from the origin of replication (2,049,646 bp from oriC), where the copy number is expected to be approximately one. We transformed this strain with the p15A plasmid encoding each DNAP of interest under the control of a rhamnose inducible promoter. Point mutations that convert the TAG stop codon to sense codons can confer chloramphenicol resistance. Second, we inoculated strains into 1 mL 2XTY media and grew them at 37 °C while shaking (220 rpm) for 12 h. Then 1 μL of the culture was inoculated into 1 mL 2XTY media supplemented with apramycin and 10 mM rhamnose, and the cultures were grown at 37 °C while shaking (220 rpm) for 12 h (∼2^10^-fold increase in cells after growth, 10 generations). All samples were then diluted and plated on plates with or without 20 µg ml^-1^ chloramphenicol, to test the proportion of chloramphenicol-resistant cells. 12 biological replicates were performed for each strain. FALCOR ^30^ was used to calculate mutation frequency (*m*) based on cell numbers on the selection and non-selection plates for all 12 biological replicates. Mutation per generation per base pair *µ* (s.p.b.) was calculated using *µ* (s.p.b.) = *m* / (R × C). For the parameter R, which is the number of distinct mutation sites that make the resistance gene effective, we found that 8/9 possible single base substitutions (which yield sense codons) can result in chloramphenicol resistance, as determined by Sanger sequencing of resistant clones. So, R = 8/3. C is the gene copy number and equals 1 because the gene is integrated at a genomic locus where the copy number is expected to be one.

To measure the *V. natriegens* genomic mutation rate, we used the endogenous genomic *rpoB* gene which can confer rifampicin resistance via single point mutations. We performed the experiments as above for *E. coli*, but supplied rifampicin at a concentration of 10 µg ml^-1^ rather than chloramphenicol. Nanopore sequencing of 95 rifampicin-resistant *V. natriegens* colonies revealed four single base substitutions that could confer rifampicin resistance, so R = 4/3. Since *rpoB* is a genomic gene, we assume C = 1 as the copy number is expected to be one.

We also used Luria-Delbrück fluctuation analysis to measure the mutation rate on the O-replicon. For *E. coli*, we created a Kan^R^-Cm^R^(Q38TAG) orthogonal replicon that contains an amber stop codon (TAG) at position 38 of the chloramphenicol resistance gene. We introduced this orthogonal replicon into the lacI6 strain. In the case of all MTAG O-DNAP variants, we transformed these cells with a p15A plasmid encoding terminal protein and a DNA polymerase of interest under the control of a rhamnose inducible promoter. In the case of all MGC O-DNAP variants, we transformed these cells with a p15A plasmid encoding terminal protein and a DNA polymerase of interest under the control of a rhamnose inducible promoter. All strains were inoculated into 1 mL 2XTY media with kanamycin, ampicillin, and grown at 37 °C while shaking (220 rpm) for 12 h. Then 1 μL of the culture was inoculated into 1 mL 2XTY media with kanamycin, ampicillin and 10 mM rhamnose, and grown at 37 °C while shaking (220 rpm) for 12 h (∼2^10^, 10 generations). All samples were then diluted and plated on plates with or without 20 µg ml^-1^ chloramphenicol, to test the proportion of chloramphenicol-resistant cells. 12 biological replicates were performed for each strain and FALCOR was used to calculate the mutation frequency (*m*). Substitutions per bp per generation, *µ* (s.p.b.), were calculated using *µ* (s.p.b.) = *m* / (R × C), where R = 8/3 and C = copy number of orthogonal replicons tested by qPCR for each sample.

For *V. natriegens*, we created a TetA-Cm^R^(Q38TAG) orthogonal replicon that contains an amber stop codon (TAG) at position 38 of the chloramphenicol resistance gene replicon and transformed this into *V. natriegens* bearing a plasmid with a RSF1010 origin encoding the synthetic replication operon and then transformed the strain with a p15A plasmid encoding a DNA polymerase of interest under the control of a constitutive promoter. All strains were inoculated into 1 ml 2XTYV media with kanamycin, tetracycline, and grown at 37 °C while shaking (220 rpm) for 10 h. All samples were then diluted and plated on plates with or without 5 µg ml^-1^ chloramphenicol, to test the proportion of chloramphenicol-resistant cells. 12 biological replicates were performed for each strain and FALCOR was used to calculate the mutation frequency (*m*). Substitutions per bp per generation, *µ* (s.p.b.), were calculated as described above.

### AlphaFold2 structure prediction

The PRD1 PolB, AdhE, AdhP, MhpF, and TetA structures were predicted using ColabFold ^31^ version of AlphaFold2 algorithm ^32^ implemented in ChimeraX (v1.5) ^33^. The structures were visualized in PyMOL (v2.5.4).

### Mutagenic DNA polymerase characterization by Illumina NGS

Polymerases were subjected to 20 generations of passaging in 3 biological replicates. The linear replicon was purified by minimal PCR amplification (Q5 polymerase, New England Biolabs), and prepared for Illumina sequencing using the NEB UltraExpress FS Library Prep Kit (E3340, New England Biolabs) and NEB Unique Dual Index UMI Adaptors (E7395, New England Biolabs) according to manufacturer’s instructions. Libraries were quantified and pooled, and sequenced on an Illumina NextSeq2000 with a P2, 100 cycle XLEAP reagent kit. NextSeq2000 generated demultiplexed fastq files with DRAGEN BCL-Convert v4.2.7. Reads were trimmed and filtered for quality (*cutadapt*) ^34^, aligned to the appropriate replicon construct (*Bowtie2*) ^35^, and filtered (*samtools view*) ^36^ to keep only unselected regions of the replicon (e.g. excluding KanR gene and 150 bp of terminal ends which are under selection). Mutational frequencies were used to calculate estimated mutation rates by dividing through 20 generations for every base mutation and indels; e.g. A to C, G, T, Ins, Dels; Ns were ignored. Since this method has no strand information, base-pair substitutions were aggregated (e.g. C to A with G to T).

Mutational spectra show the mean of positive mutation rates for each substitution mutation, and the overall DNAP mutation rate was taken as the mean of all the summed median mutation substitution rates of A, C, G and T.

### Accelerated continuous evolution of an ethanol assimilation pathway in *E. coli*

We first created an O-replicon harboring the *adhE-adhP-mhpF* operon under the control of a constitutive PM4 promoter and transformed this into the lacI6 strain and then transformed the strain with a p15A plasmid encoding a mutagenic O-DNAP variant MTAG1 under the control of a salicylic acid inducible promoter. For the continuous evolution experiment, we performed three replicates. We inoculated all strains into 5 mL media with 10 µM salicylic acid (to induce O-replicon mutagenesis mediated by the MTAG1 O-DNAP) in 50 mL falcon tubes and grew these cultues at 37 °C while shaking (220 rpm) for 12 h. Then we passaged cells harbouring this replicon in 50 mL M9 minimal medium supplemented with 10% (v/v) LB, 10 µM salicylic acid, 20 g L^−1^ ethanol, and decreasing concentrations of glucose. After five passages, we obtained pools of cells that grew to much higher maximum cell densities relative to the input control. We then cloned different hits from three independent evolution replicates onto a standard circular plasmid (colE1) for validation.

### Accelerated continuous evolution of *tetA* in *V. natriegens*

We first created a TetA-Cm^R^(Q38TAG) orthogonal replicon and transformed this into *V. natriegens* bearing a plasmid with a RSF1010 origin encoding the synthetic replication operon and then transformed the strain with a p15A plasmid encoding a mutagenic O-DNAP variant VOD6 under the control of a constitutive promoter. For the continuous evolution experiment, we performed 4 replicates. We inoculated all strains into 5 mL 2XTYV (2XTY with 10 x V2 salts) media in 50 mL falcon tubes and grew them at 37 °C while shaking (220 rpm) for 7 h. Then, 500 μL of the culture was inoculated into 50 mL 2XTYV media with kanamycin, tetracycline, 0.1 mM IPTG, and 0.1 µg mL^-1^ tigecycline. We passaged cells once all replicates had reached OD_600_ > 1 and diluted the culture into 50 mL 2XTYV media with kanamycin, tetracycline, 0.1 mM IPTG, and increasing concentrations of tigecycline (from 0.25 µg mL^-1^ to 30 µg mL^-1^). We completed 5 passages in 16 h. After the final passage under continuous mutagenesis, we passaged cells into rhamnose-free medium to switch to using the WT O-DNAP, thereby switching off further mutagenesis. The following day, we cloned the evolved sequences from each of the four replicates into a p15A circular plasmid backbone in *E. coli*. The pooled plasmids were then transformed back into *V. natriegens* to ensure that each cell contained a single clonal sequence and that the plasmid copy number was consistent across all mutants.

## Supporting information

All Supplementary Information

## Acknowledgments

We thank the FACS facility (MRC-LMB) (Pier Andrée Penttilä and Fan Zhang) for support.

## Author contributions

J.W.C., R.T. and F.B.H.R. conceptualized the project. R.T. optimized the electroporation protocol for establishing O-replicons, designed and optimized approaches to engineer the O-replicon *in vivo*, established the miniprep and retransformation method of the O-replicon, designed the replicon-REXER. R.T. and M.K. performed the replicon-REXER experiments. R.T., K.J. and P.S.Z. did the experiments on evolving the O-DNAP in *E.coli*. R.T. and M.K. performed the accelerated continuous evolution of ethanol assimilation pathway. F.B.H.R. established and investigated the stability of the O-replication system in *V. natriegens* and *P. putida*. R.T. evolved the O-DNAP in *V. natriegens*. R.T. and F.B.H.R. performed the accelerated continuous evolution of *tetA* in *V. natriegens*. M.K. analyzed structures of evolved proteins obtained from the continuous evolution experiments. P.S.Z. optimized the transformation protocol for *V. natriegens*. K.C.L. did NGS analysis. J.W.C., F.B.H.R. and R.T. wrote the paper with input from all authors. J.W.C. supervised the project.

## Data and materials availability

All data are available in the main text or supplementary materials. The computer code to analyze NGS data is available from Zenodo (https://doi.org/10.5281/zenodo.10213374). The authors agree to provide any materials and strains used in this study upon request.

## Supplementary Materials

Supplementary Figs. 1 to 34

Extended Data Table 1

