## Supplementary material for "A portable orthogonal replication system enables continuous gene evolution near the biological speed limit": All Supplementary Information

**The PDF file includes:**

Supplementary Figs. 1 to 34

Extended Data Table 1

References (*1-6*)

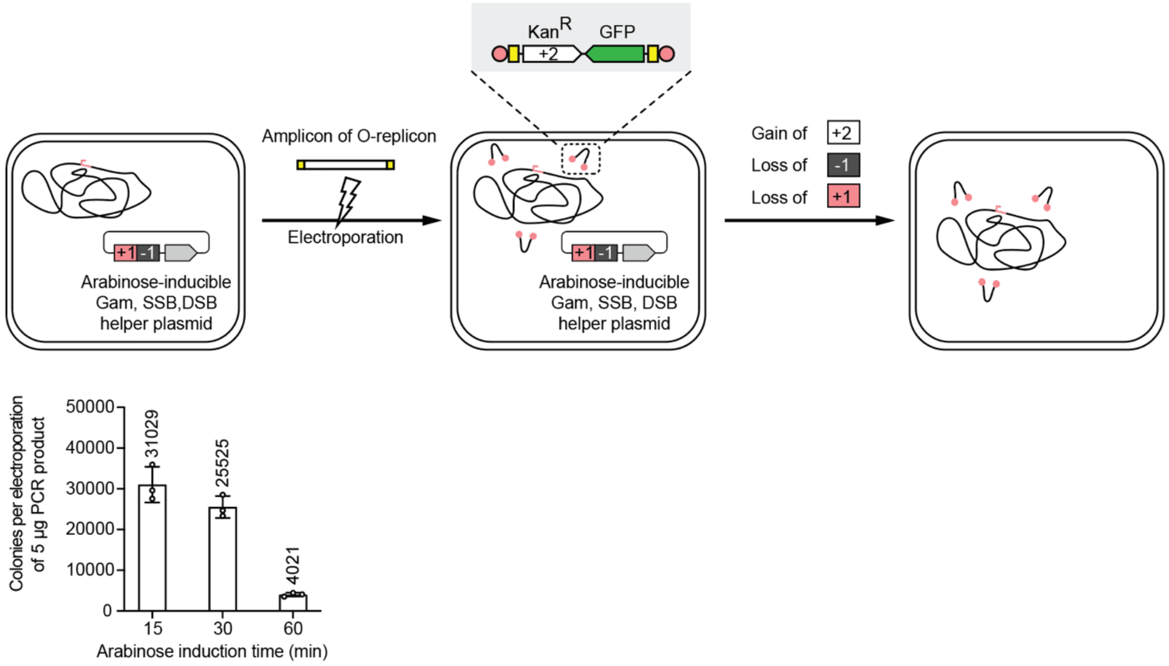


**Supplementary Figure 1. Increasing the efficiency of O-replicon establishment from amplicons by optimizing the induction time of the helper plasmid.**

The length of the replicon is 2.2 kb. The increase in efficiency was achieved by shortening the induction time of the helper plasmid from 60 to 15 minutes. After the replicon is established, the helper plasmid is readily cured by selection for loss of the -1 negative selection marker. n = 3, data are shown as mean ± s.d..

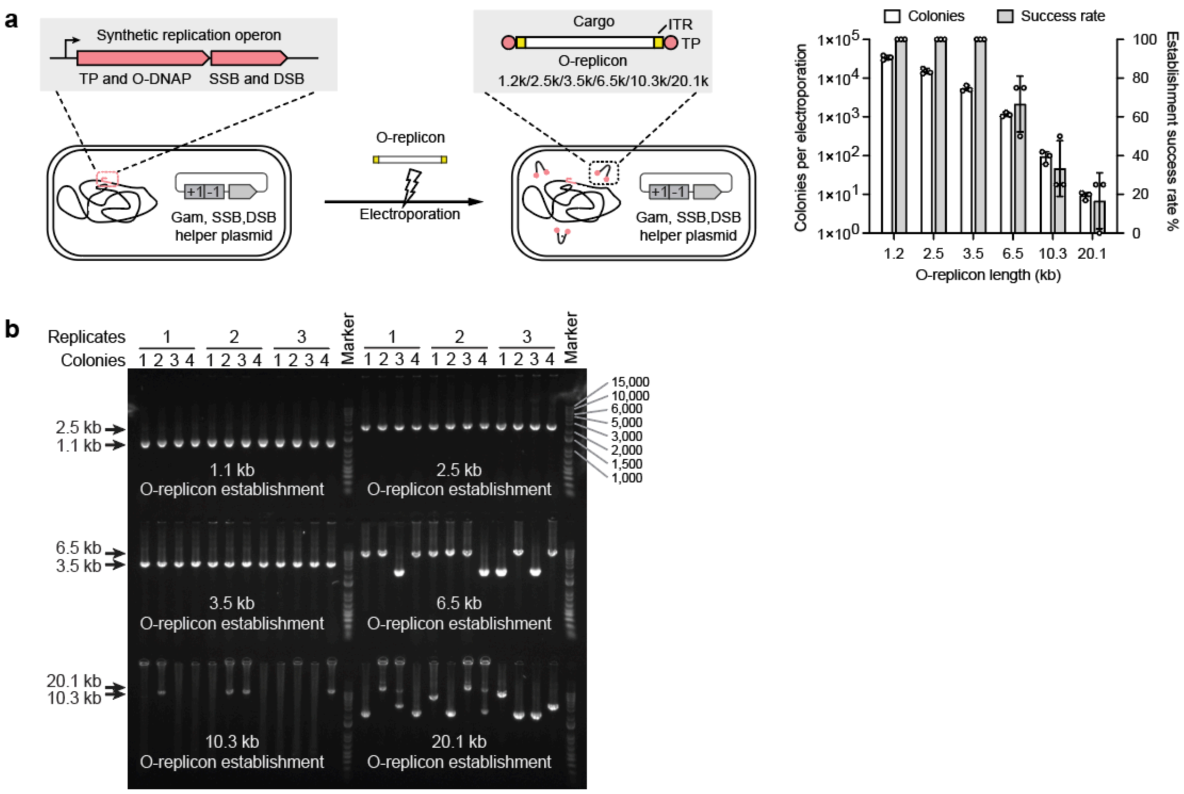


**Supplementary Figure 2. Increased efficiency of establishing O-replicons from amplicons enables the establishment of varying lengths of replicons.**

**a**, ITR-flanked amplicons of different lengths were electroporated into cells harboring a genomic synthetic replication operon and a helper plasmid. n = 3, error bars are ± s.d. TP, terminal protein; O-DNAP, orthogonal DNA polymerase; SSB, single-stranded DNA binding protein; DSB, double-stranded DNA binding protein; ITR, inverted terminal repeat (in yellow). **b**, Genotyping establishment of the O-replicons of varying lengths. The lengths of the resultant O-replicons were verified by colony PCR and the amplicon sequences were verified via Nanopore sequencing.

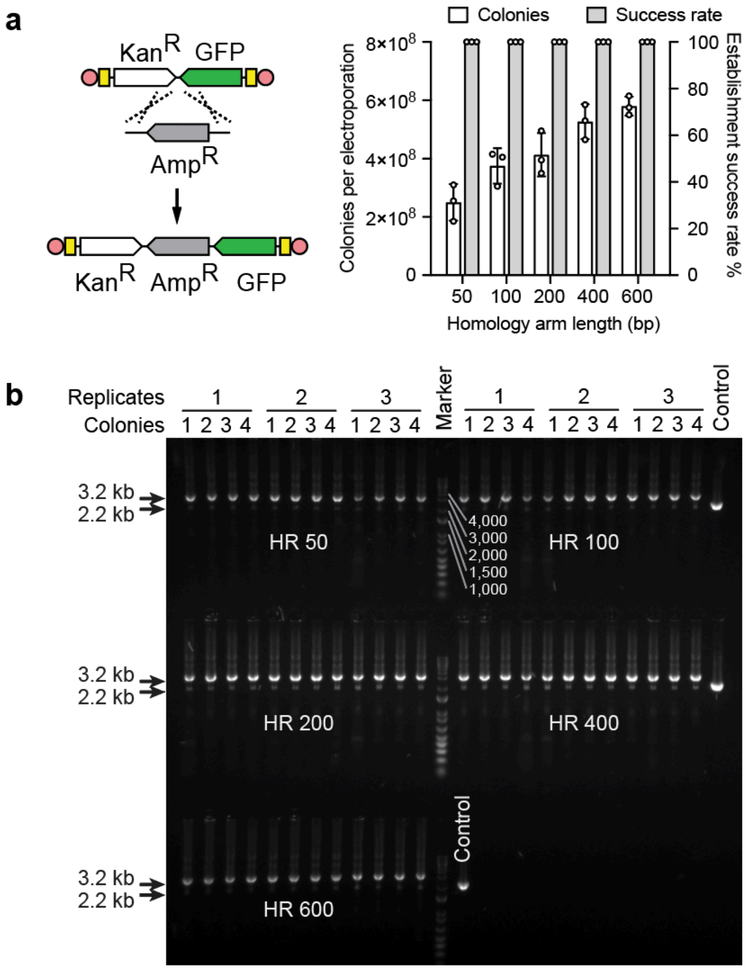


**Supplementary Figure 3. Engineering an established O-replicon using λ Red recombination.**

**a**, Donor DNA amplicons flanked by varying lengths of homology arms (HR) were electroporated into cells harboring a genomic synthetic replication operon, a Kan$^R$-GFP replicon, and a λ Red plasmid (n = 3, data are shown as mean± s.d.). **b**, Genotyping gene insertion into O-replicons. The length of the O-replicons was verified by colony PCR and the amplicon sequences were verified via Nanopore sequencing. The control is an amplicon of the O-replicon before λ Red-mediated insertion of the *ampR* gene.

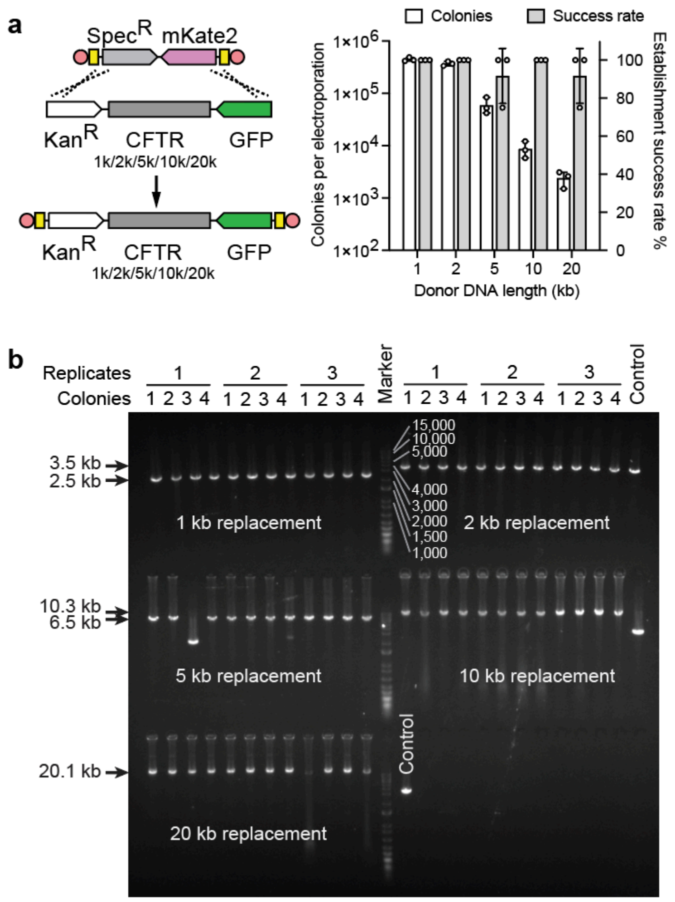


**Supplementary Figure 4. λ Red recombination enables reliable establishment of new replicons of varying lengths.**

**a**, 600 bp homology arms (HR)-flanked amplicons of different lengths were electroporated into cells harboring a genomic synthetic replication operon, a Spec$^{R}$-mKate2 replicon, and a λ Red plasmid. CFTR, DNA sequences from the human CFTR gene. n = 3, data are shown as mean ± s.d. **b**, Genotyping gene replacement of the O-replicons. The length of the O-replicons was verified by colony PCR and the amplicon sequences were verified via Nanopore sequencing. The control is the amplicon of the O-replicon before λ Red engineering.

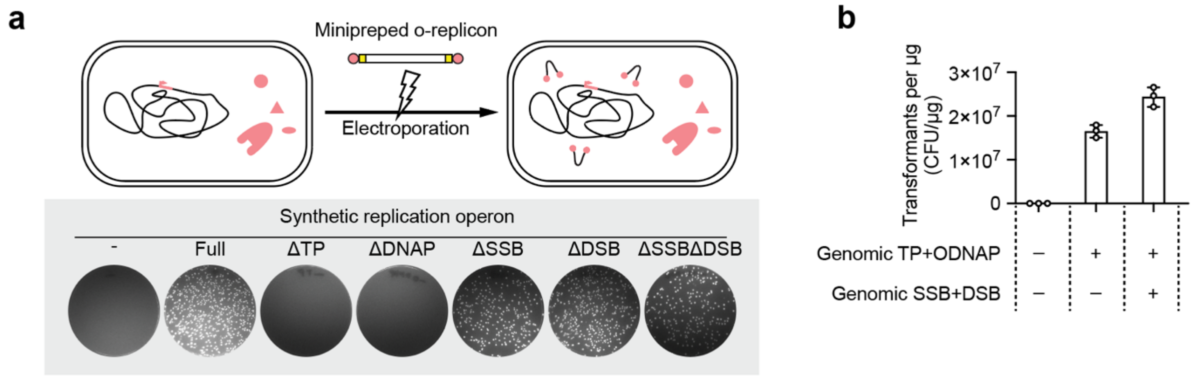


**Supplementary Figure 5. Transformation of extracted O-replicons reveals the minimal synthetic replication operon.**

**a**, Shown are plates of extracted O-replicons into strains with the indicated disruptions to the genes in the synthetic replication operon. **b**, Extracted replicons can be efficiently transformed into cells. TP, terminal protein; O-DNAP, orthogonal DNA polymerase; SSB, single-stranded DNA binding protein; DSB, double-stranded DNA binding protein. n = 3, data are shown as mean ± s.d.

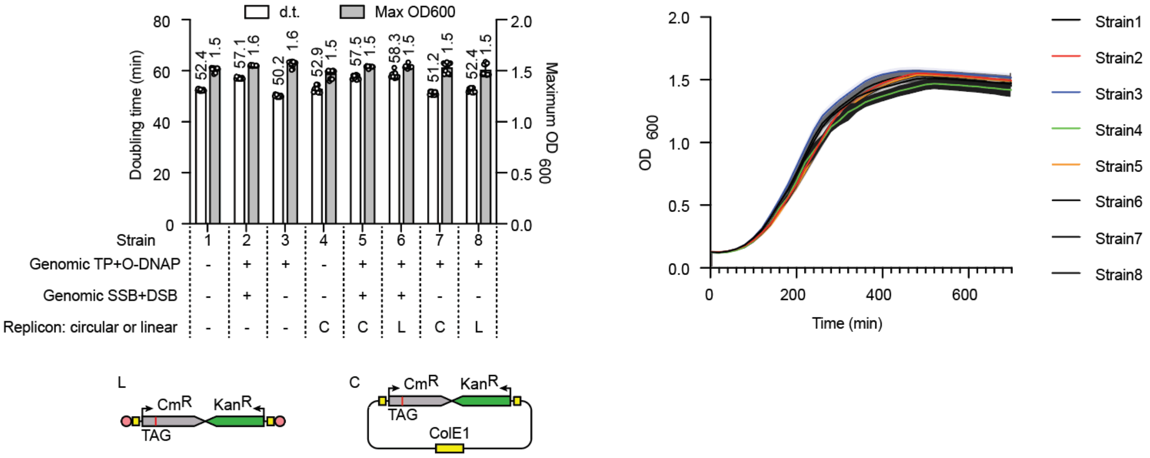


**Supplementary Figure 6. Testing growth of wild-type *E. coli* and *E. coli* strains harboring different genomic synthetic replication operons and different replicons.** L, a Kan$^R$-Cm$^R$(Q38TAG) linear O-replicon, that contains an amber stop codon (TAG) at position 38 of the chloramphenicol resistance gene (Cm$^R$). C, a circular ColE1 plasmid with identical sequence to the orthogonal replicon. D.t., doubling time. n = 12, data are shown as mean± s.d..

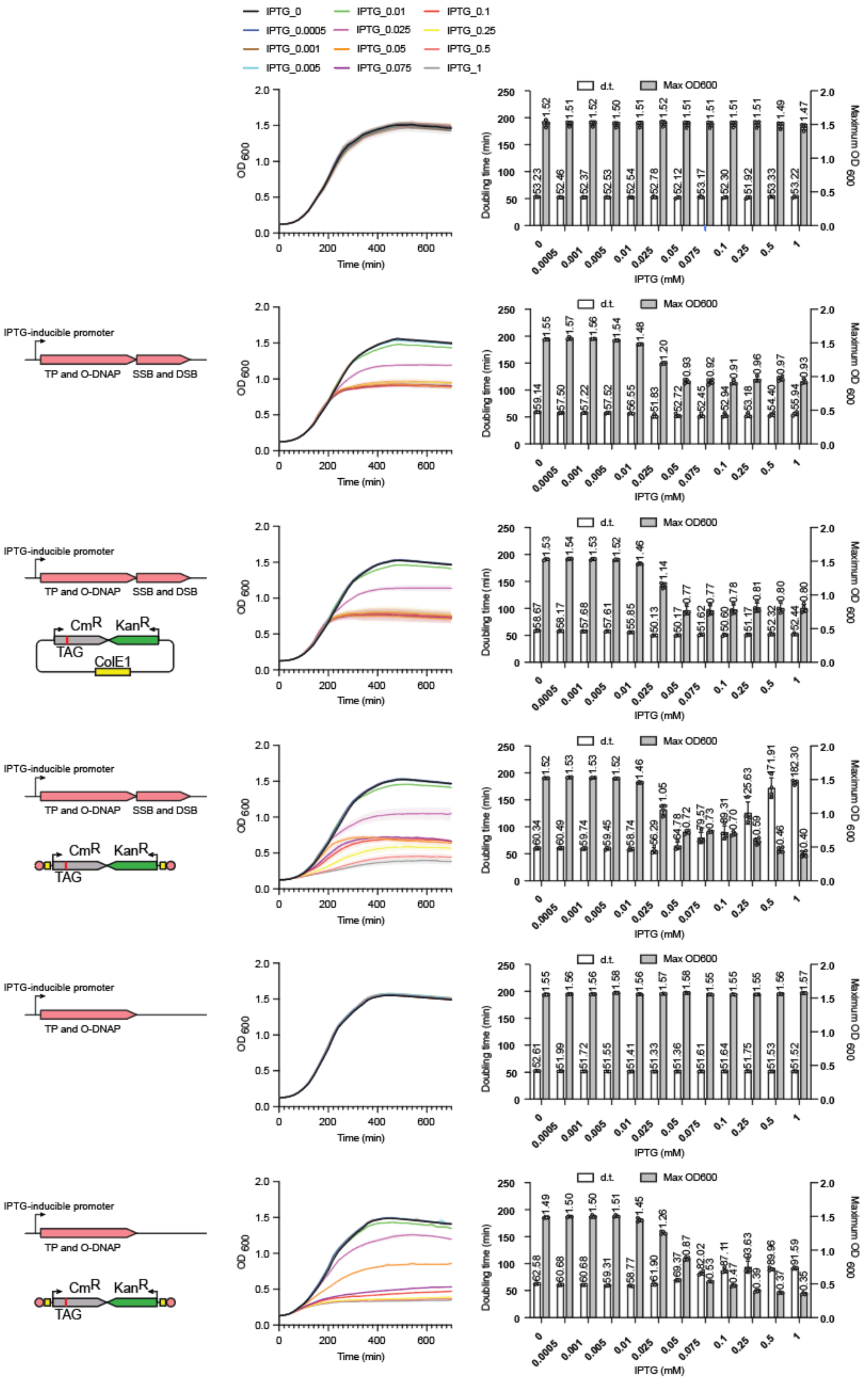


**Supplementary Figure 7. Testing growth of wild-type *E. coli* and *E. coli* strains harboring different genomic synthetic replication operons and different replicons under different concentrations of IPTG induction.** D.t., doubling time. n = 4, data are shown as mean ± s.d.

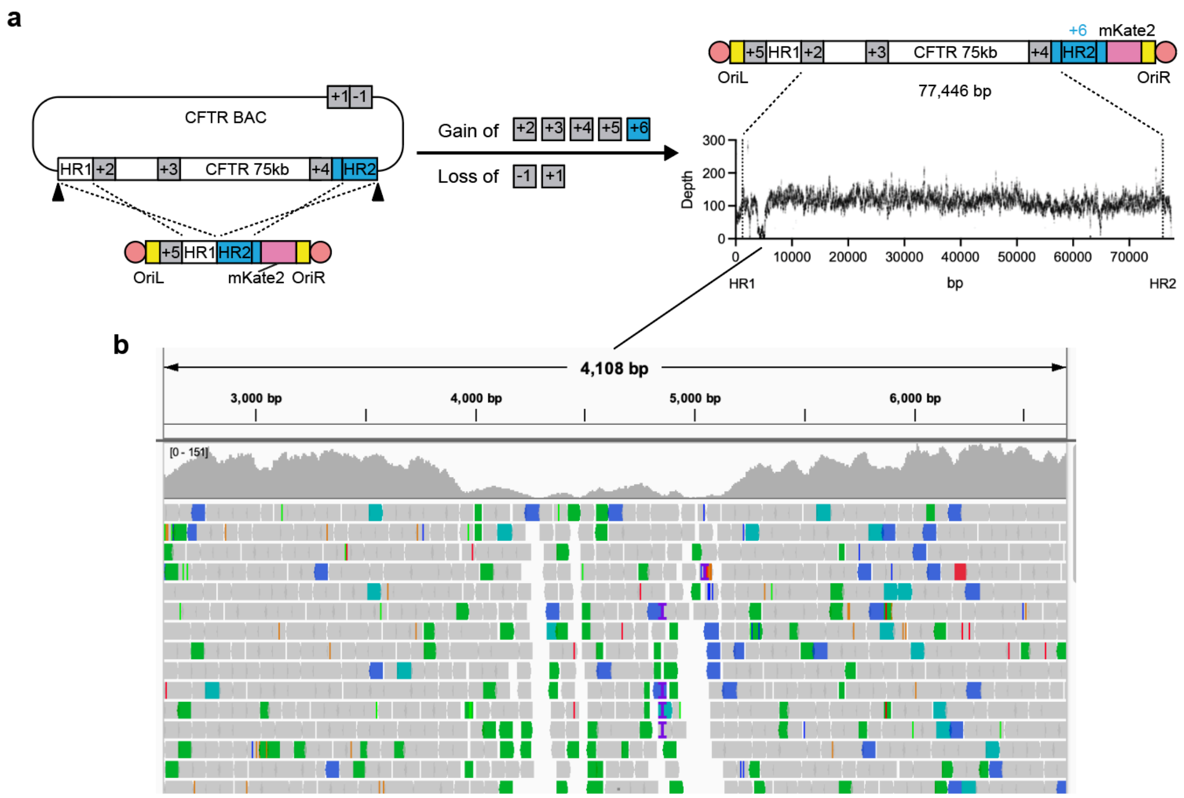


**Supplementary Figure 8. Replicon-REXER enables the establishment of a 77 kb replicon.**

**a**, Schematic of Replicon-REXER. +1, *zeoR*; +2, *kanR*; +3, *gmR*; +4, ampR; +5, *specR*; -1, *rpsL*; Homology arm 1 (HR1), 90 bp; HR2, 300 bp. An intact chloramphenicol-resistance gene (+6) only upon successful recombination. Shown is Illumina sequencing read coverage. **b**, Screenshot of the Illumina sequencing results for the 1058 bp sequence with 76% GC content, exhibiting a low read depth, and the surrounding sequence with higher read depth.

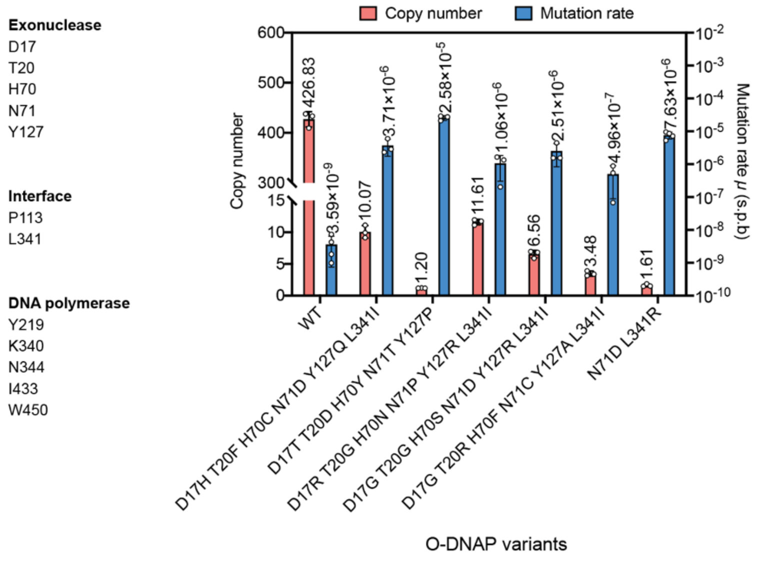


**Supplementary Figure 9. Directed evolution of mutagenic O-DNAPs by site-saturation mutagenesis.**

The mutation rates were measured after 10 generations via fluctuation tests. For assessment of the orthogonal replicon mutation rate, we used an O-replicon-encoded *Cm*$^R$ gene with a TAG stop codon at position 38 and each O-DNAP variant was expressed from a p15A plasmid via rhamnose induction (5 mM). n = 3, data are shown as mean ± s.d.

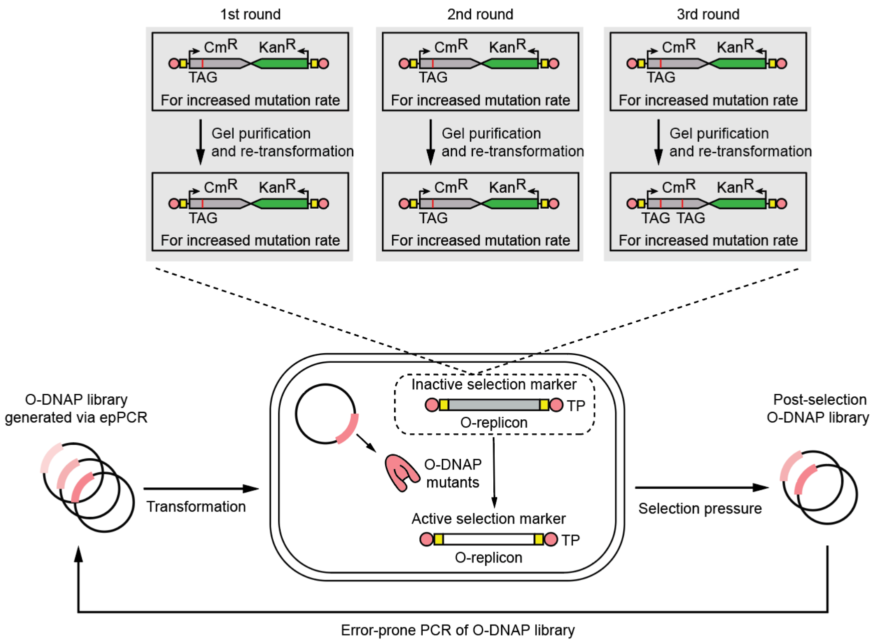


**Supplementary Figure 10. Overview of the directed evolution of mutagenic O-DNAPs via the TAG-inactivated Cm$^R$ gene.** O-DNAP libraries were generated using error-prone PCR. We did three rounds of directed evolution with the purpose of increasing the mutation rate. For the first round of directed evolution, we first generated an O-DNAP library using error-prone PCR targeting the DNA polymerase domain using the rationally designed Y127A O-DNAP mutant as a template. The library was then transformed into cells harboring a Cm$^R$-Kan$^R$ reporter O-replicon where the Cm$^R$ gene was inactivated via the introduction of a TAG stop codon. We then passaged the resultant transformants for 10 generations to allow for the O-replicon mutagenesis to occur. Reversion of the stop codon to a sense codon via mutations introduced by a mutagenic O-DNAP culminate in a functional Cm$^R$ gene that confers chloramphenicol resistance to the cell. Cells were then selected using chloramphenicol, and the O-DNAP plasmid library was extracted from these cells. Then the selected library was gel purified to remove the mutated replicon from the last selection and transformed into cells harboring the same unmutated reporter O-replicon for the next selection step. After two rounds of enrichment, we generated a new library for the second round using error-prone PCR targeting the DNA polymerase domain using the enriched library from the first round as a template. After two further rounds of enrichment, we generated a new library for the third round using error-prone PCR targeting both the exonuclease domain and the DNA polymerase domain using the enriched library from the second round as a template. The final selection steps were conducted using a doubly TAG-inactived Cm$^R$ reporter O-replicon.

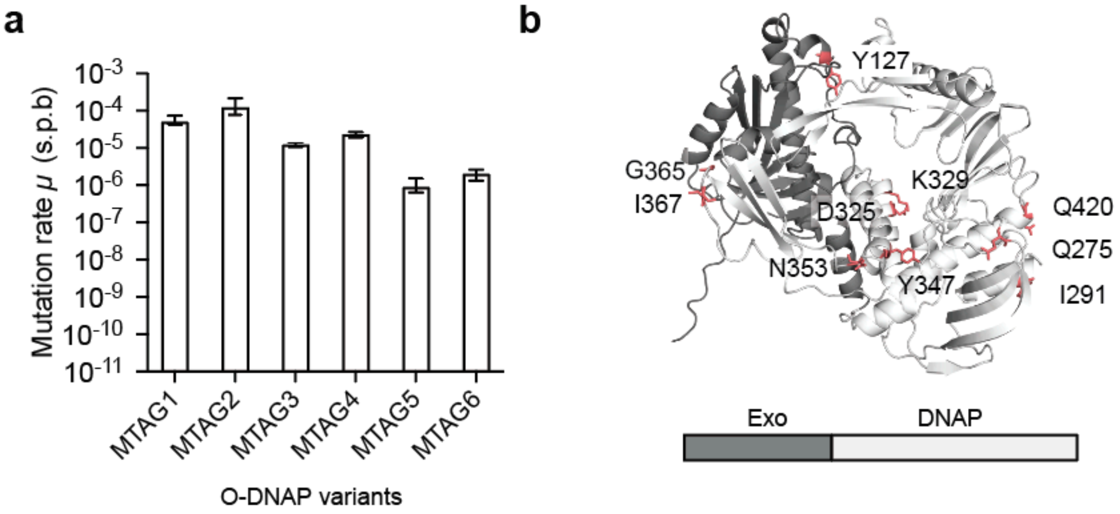


**Supplementary Figure 11. O-DNAP variants evolved via the TAG-inactivated Cm$^R$ gene.**

**a**, Determination of orthogonal replicon mutation rate ($\mu$, s.p.b.) for O-DNAP variants. The mutation rate was measured after 10 generations via fluctuation tests. For assessment of the orthogonal replicon mutation rate, we used an orthogonal replicon-encoded *Cm$^R$* gene with a TAG stop codon at position 38 and the O-DNAP variants were expressed from a p15A plasmid via rhamnose induction (5 mM). The identical data for MTAG1 and MTAG2 are shown in Fig.1C. **b**, AlphaFold2-predicted model of the PRD1 DNAP. Residues enriched in the error-prone O-DNAP variants are shown in stick representation, labelled and highlighted in red.

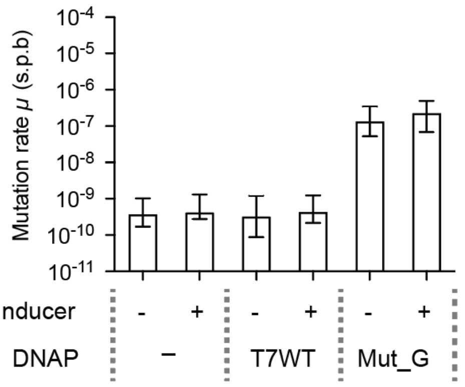


**Supplementary Figure 12. The expression of an error-prone T7 DNAP variant resulted in a high level of genome mutagenesis.** The WT T7 DNAP or an error-prone variant (Mut_G) were expressed from a p15A plasmid under the control of a salicylic acid-inducible promoter. For the induced samples, 10 µM salicylic acid was added. The mutation rates were determined by fluctuation tests which measured the reversion of a TAG stop codon in a genomically-integrated chloramphenicol resistance gene. For all experiments n = 12, data are shown as median ± upper/lower 95% bounds.

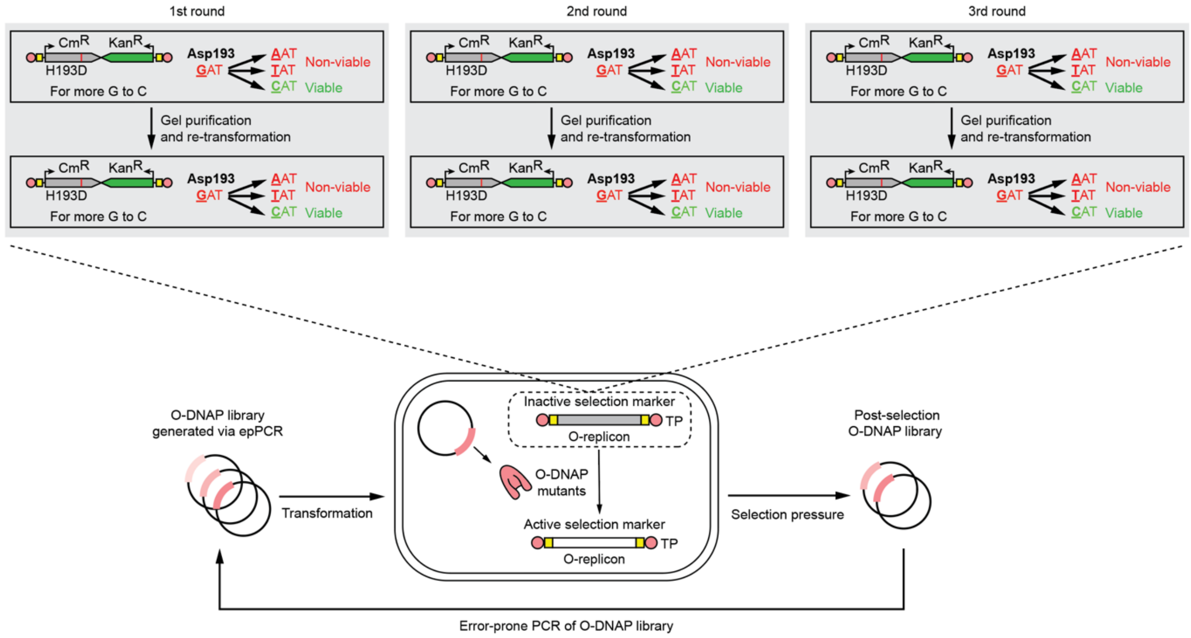


**Supplementary Figure 13. Overview of the directed evolution of mutagenic O-DNAPs via the H193D-inactivated $Cm^R$ gene.**

O-DNAP libraries were generated using error-prone PCR. We did three rounds of directed evolution with the purpose of improving the mutational spectrum. For the first round of directed evolution, we first generated an O-DNAP library using error-prone PCR targeting both the exonuclease domain and the DNA polymerase domain using the rationally designed Y127A O-DNAP mutant as a template. The library was then transformed into cells harboring a $Cm^R$-$Kan^R$ reporter O-replicon where the $Cm^R$ gene was inactivated via mutation of the essential His193 residue to Asp. We then passaged the resultant transformants for 10 generations to allow for the O-replicon mutagenesis to occur. Reversion of the Asp193 to His via mutations introduced by a mutagenic O-DNAP culminate in a functional $Cm^R$ gene that confers chloramphenicol resistance to the cell. Cells were then selected using chloramphenicol, and the O-DNAP plasmid library was extracted from these cells. Then the selected library was gel purified to remove the mutated replicon from the last selection and transformed into cells harboring the same unmutated reporter O-replicon for the next selection step. After two rounds of enrichment, we generated a new library for the second round using error-prone PCR targeting both the exonuclease domain and the DNA polymerase domain using the enriched library from the first round as a template. After two rounds of enrichment, we generated a new library for the third round using error-prone PCR targeting both the exonuclease domain and the DNA polymerase domain using the enriched library from the second round as a template.

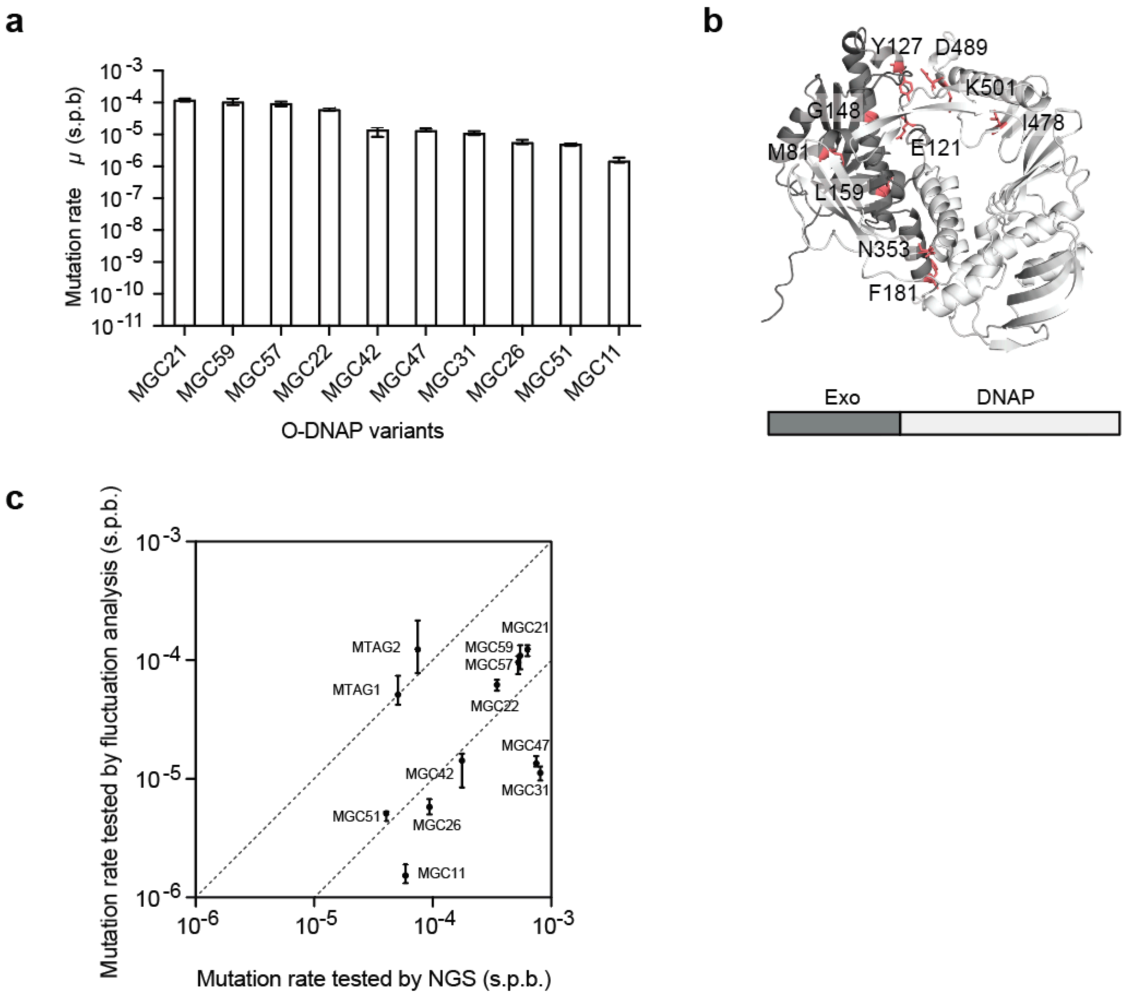


**Supplementary Figure 14. O-DNAP variants evolved via the H193D-inactivated Cm$^R$ gene.**

**a**, Determination of orthogonal replicon mutation rate ($\mu$, s.p.b.) for O-DNAP variants. The mutation rates were measured after 10 generations via fluctuation tests. For assessment of the orthogonal replicon mutation rate, we used an orthogonal replicon-encoded $Cm^R$ gene with a TAG stop codon at position 38 and the O-DNAP variants were expressed from a p15A plasmid via rhamnose induction (5 mM). The identical data for MGC26 and MGC59 are shown in Fig. 1C. **b**, AlphaFold2-predicted model of PRD1 DNAP. Residues enriched in the error-prone O-DNA polymerase variants are shown in stick representation, labelled and highlighted in red. **c**, Correlation between mutation rates tested by fluctuation analysis and NGS.The identical mutation rate data by NGS analysis are shown in Fig. 1e. The identical mutation rate data by fluctuation analysis are shown in Fig. 1c, fig. S11, and **a**.

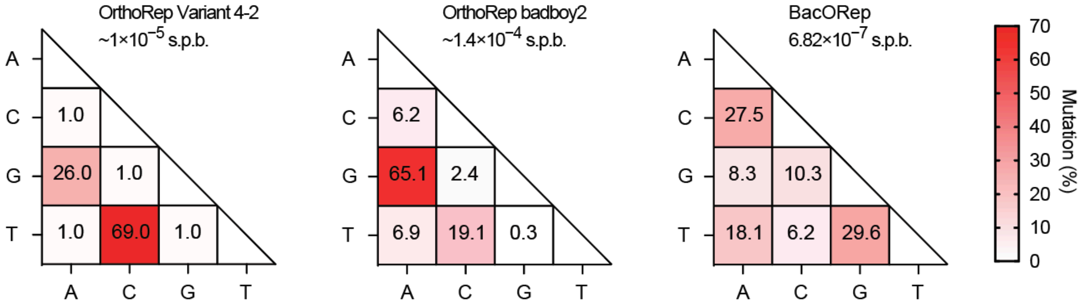


**Supplementary Figure 15. Mutational spectra for three mutagenic DNAPs used in orthogonal replication systems [1–3] .**

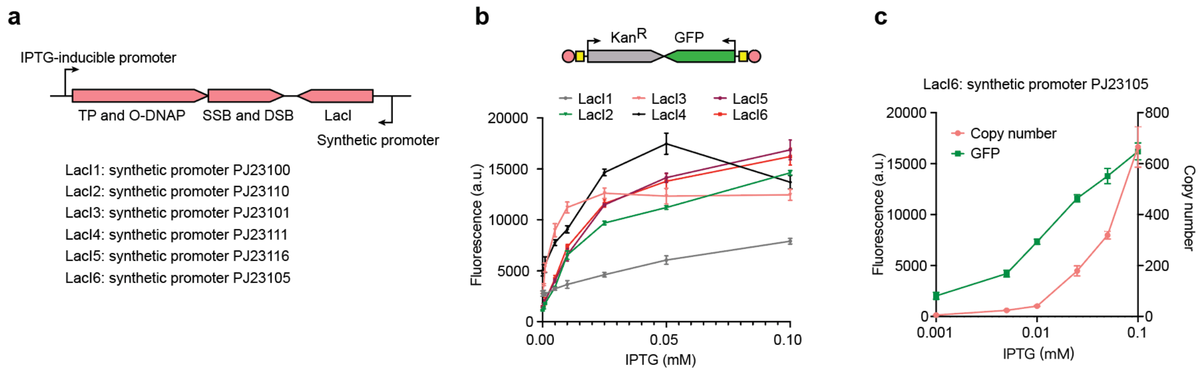


**Supplementary Figure 16. Control of orthogonal replicon copy number range. a**, Control of the orthogonal replicon copy number range is achieved by controlling the expression level of LacI using different synthetic promoters [4]. **b**, GFP fluorescence (normalized to $OD_{600}$) at different IPTG concentrations (n=3, data are shown as mean ± s.d.). **c**, Correlation between $Kan^R$-GFP orthogonal replicon copy number and GFP fluorescence (n=3, data are shown as mean ± s.d.). Orthogonal replicon copy number, as determined by qPCR, and GFP fluorescence (normalized to $OD_{600}$) of LacI6 were measured at different IPTG concentrations. GFP fluorescence data from **b** was replotted for **c**.

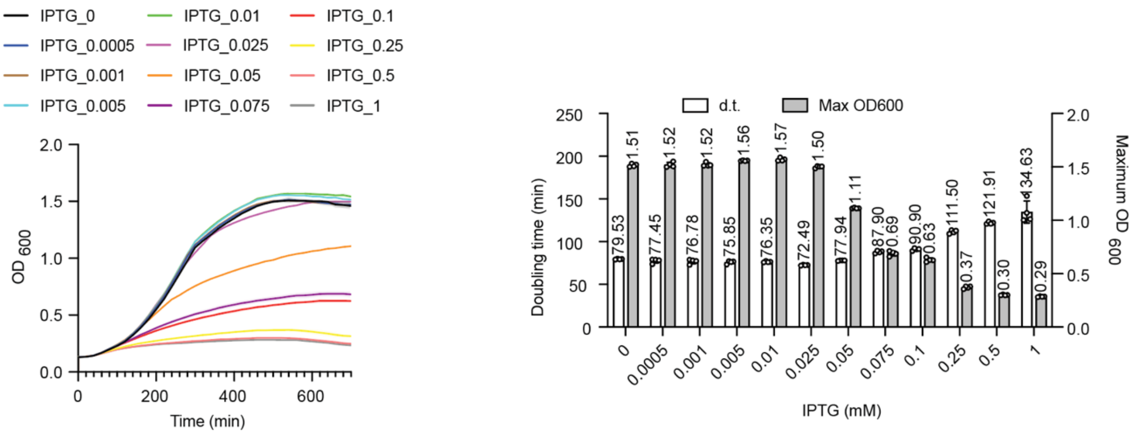


**Supplementary Figure 17. Testing growth of the LacI6 strain harboring an O-replicon under different concentrations of IPTG induction** (n = 4, data are shown as mean ± s.d.)**.** The O-replicon is a $Kan^R$-$Cm^R$(Q38TAG) orthogonal replicon, that contains an amber stop codon (TAG) at position 38 of the chloramphenicol resistance gene ($Cm^R$). d.t., doubling time.

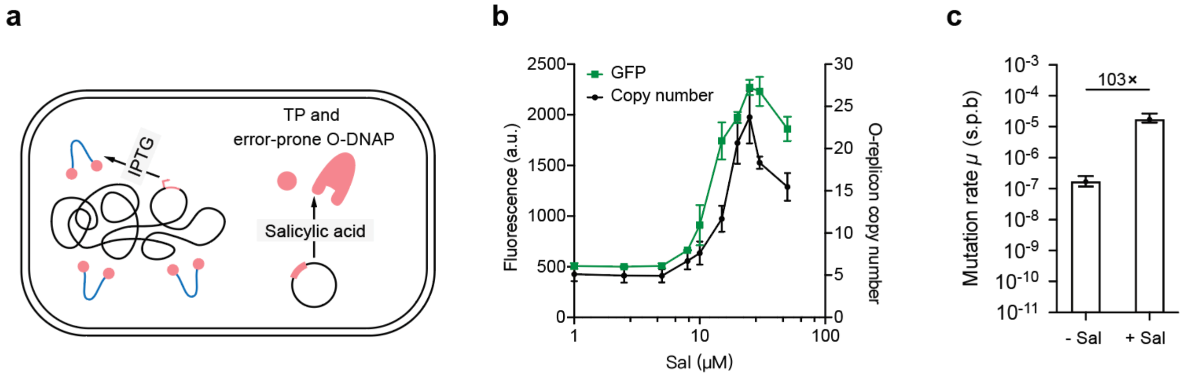


**Supplementary Figure 18. Inducible error-rate via controlling the expression of TP and error-prone O-DNAP from a plasmid.**

**a**, The *E. coli* strain for continuous evolution experiment is a LacI6 strain harboring a genomic synthetic replication operon and a pSC101 plasmid encodes for TP and the O-DNAP variant MTAG1 under the control of a salicylic acid-inducible promoter. Control of orthogonal replicon copy number can be achieved by inducing expression of the genomic TP and WT O-DNAP via IPTG addition, or by inducing expression of the plasmid-encoded TP and a O-DNAP variant MTAG1 via salicylic acid addition. **b**, O-replicon copy number and GFP fluorescence (normalized to $OD_{600}$) with different combinations of salicylic acid (Sal) concentrations. The O-replicon copy number was determined by qPCR (n = 3, data are shown as mean± s.d.). **c**, O-replicon mutation rate with or without induction. The mutation rate was measured after 10 generations with fluctuation tests. For assessment of the O-replicon mutation rate, we used an O-replicon-encoded $Cm^R$ gene with a TAG stop codon at position 38. n = 12, data are shown as median ± upper/lower 95% bounds.

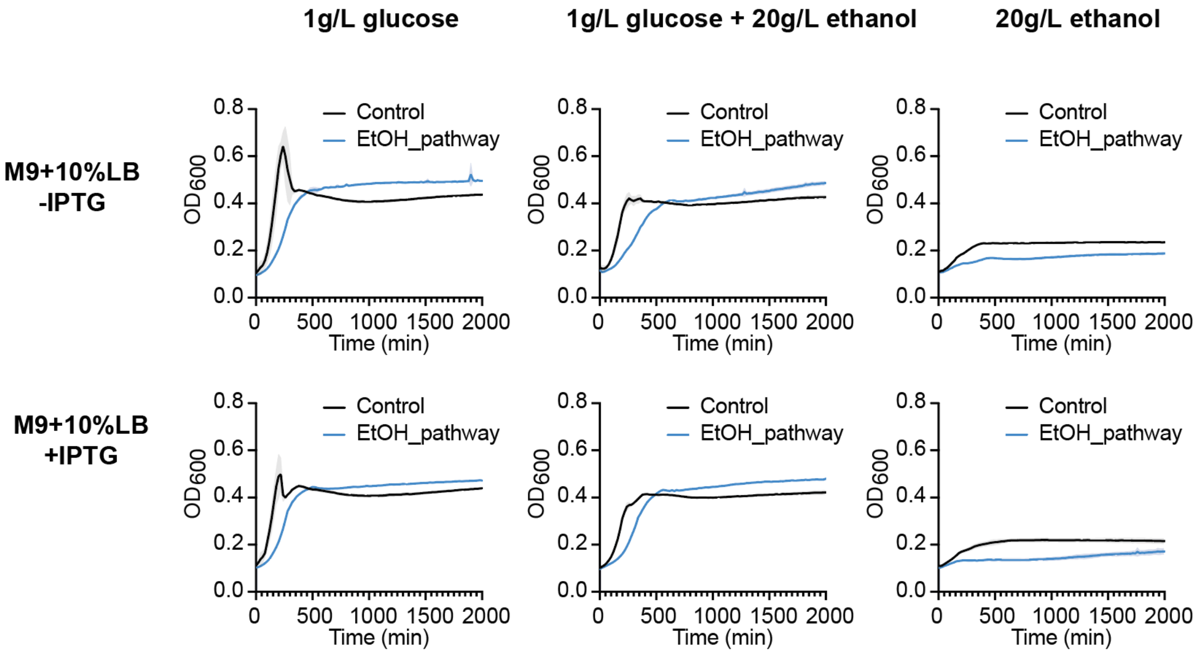


**Supplementary Figure 19. Testing growth of the LacI6 strain or the strain harboring an O-replicon encoding an ethanol assimilation pathway in different media** (n = 4, data are shown as mean ± s.d.)**.**

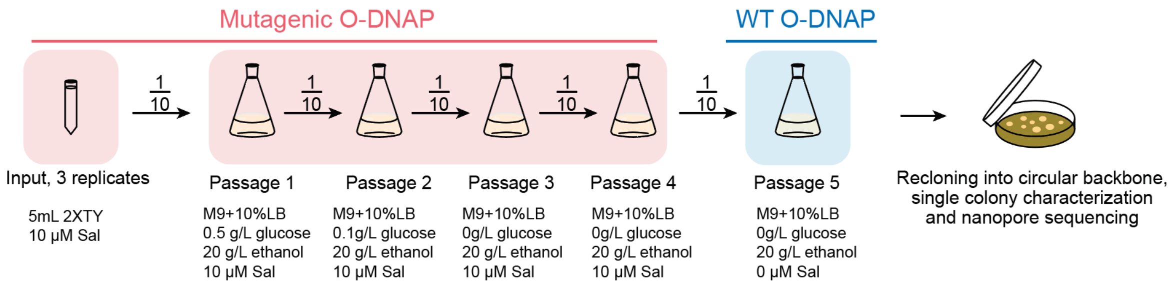


**Supplementary Figure 20. Overview of the ethanol assimilation pathway accelerated continuous evolution. Sal, salicylic acid**

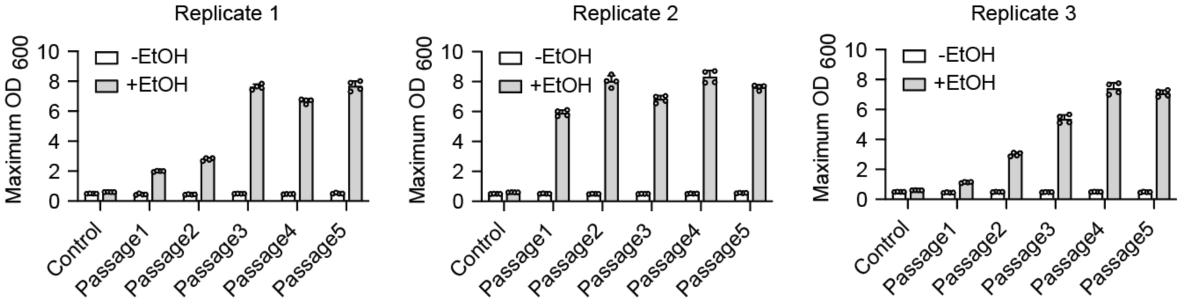


**Supplementary Figure 21. Maximum $OD_{600}$ of three evolved pools carrying the pathway-encoded orthogonal replicon after passages.** The test was done in 24 well plates (n = 4, data are shown as mean ± s.d.). We note that the measurements are in technical repeats. The control is the wild-type ethanol assimilation pathway before evolution.

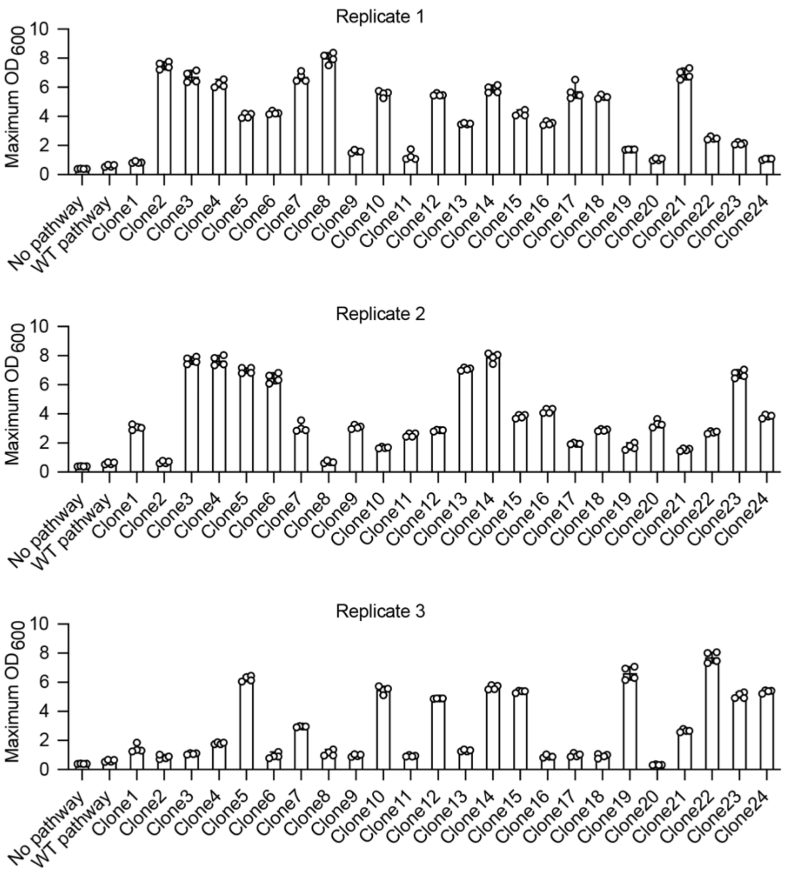


**Supplementary Figure 22. Testing maximum $OD_{600}$ of strains harboring evolved pathways on a pSC101 plasmid.** The test was done in 24 well plates (n = 4 technical replicates, data are shown as mean ± s.d.).

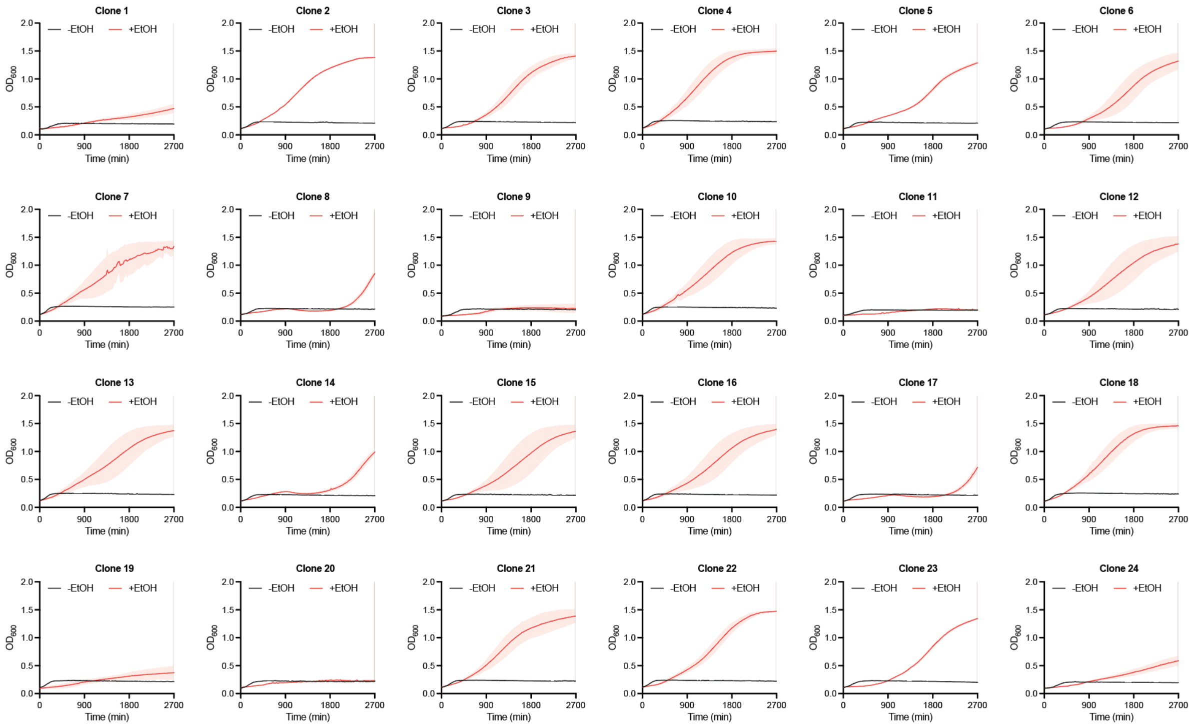


**Supplementary Figure 23. Testing growth of strains harboring evolved pathways from replicate one on a pSC101 plasmid.** The test was done in 96 well plates (n = 4, data are shown as mean ± s.d.).

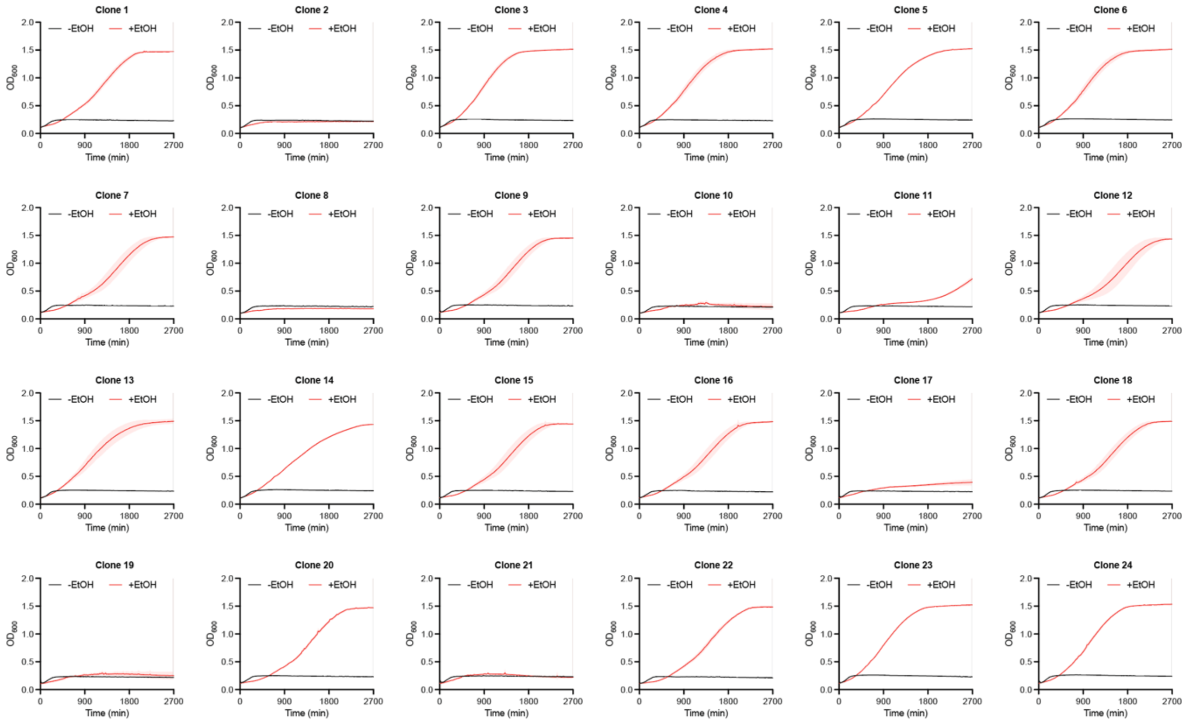


**Supplementary Figure 24. Testing growth of strains harboring evolved pathways from replicate two on a pSC101 plasmid.** The test was done in 96 well plates (n = 4, data are shown as mean ± s.d.).

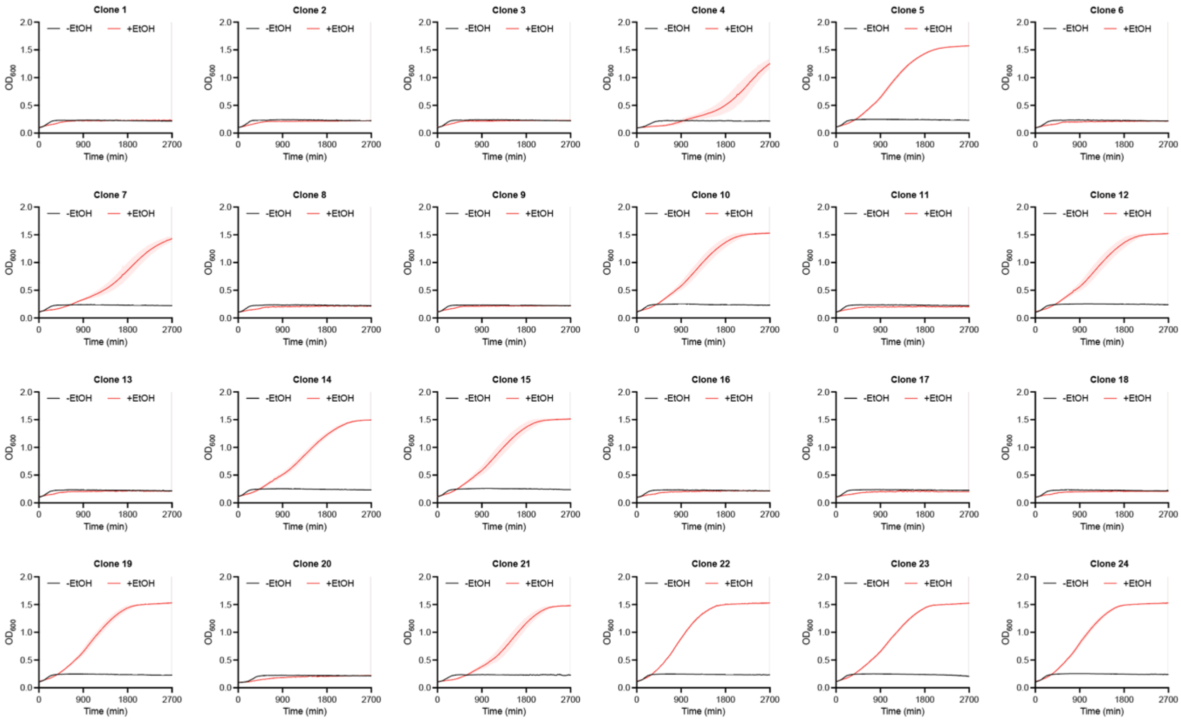


**Supplementary Figure 25. Testing growth of strains harboring evolved pathways from replicate three on a pSC101 plasmid.** The test was done in 96 well plates (n = 4, data are shown as mean ± s.d.).

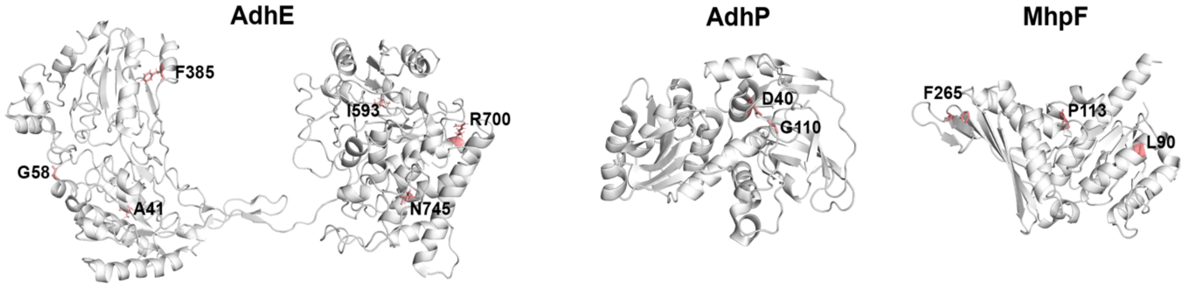


**Supplementary Figure 26. AlphaFold2 models of AdhE, AdhP, and MhpF.** The mutations that enriched within replicates in the continuous evolution experiment are shown in stick representation, labelled and highlighted in red.

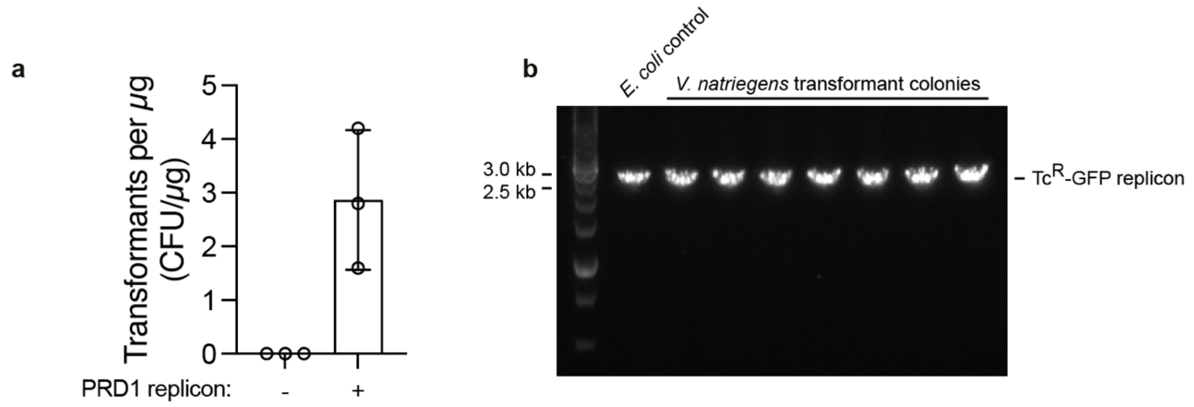


**Supplementary Figure 27. Establishing O-replicons in *V. natriegens*. a**, Efficiency of establishing orthogonal replicons in *V. natriegens* by chemically competent transformation of a $Tc^R$-GFP extracted O-replicon. n = 3, data are shown as mean ± s.d. **b**,Genotyping establishment of the O-replicons in *V. natriegens*.

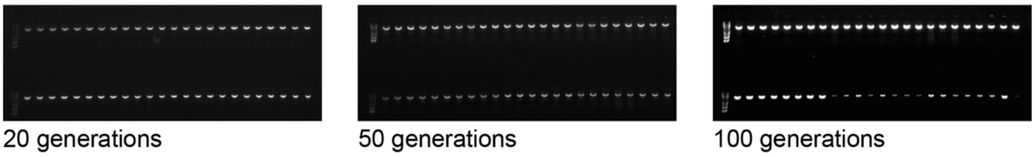


**Supplementary Figure 28. Stability of O-replicons in *V. natriegens*.** After multiple generations of passaging, 12 colonies were picked from each replicate for genotyping to verify the presence of the O-replicon. All colonies retained the replicon after 100 generations (n = 4).

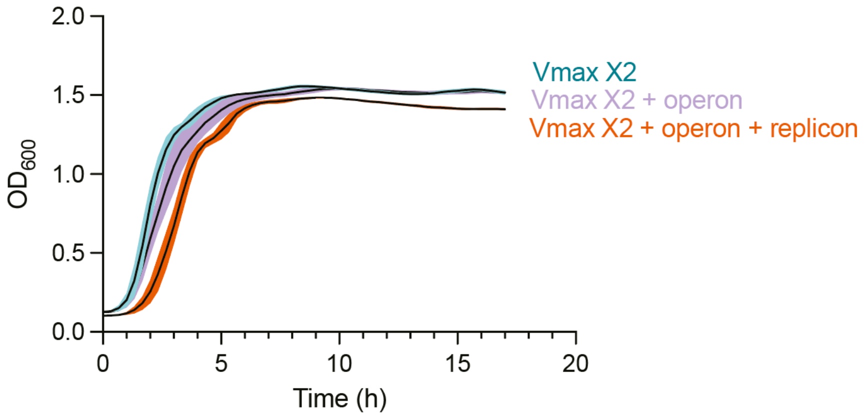


**Supplementary Figure 29.** Growth curves of wild-type *V. natriegens* and *V. natriegens* harboring a replicon (n = 8, data are shown as mean ± s.d.). The synthetic replication operon was under the control of an IPTG inducible promoter (PtacIPTG). The Kan$^R$-RFP replicon consists of flanking 110 bp ITR, a kanamycin resistance gene and an RFP gene (mKate2).

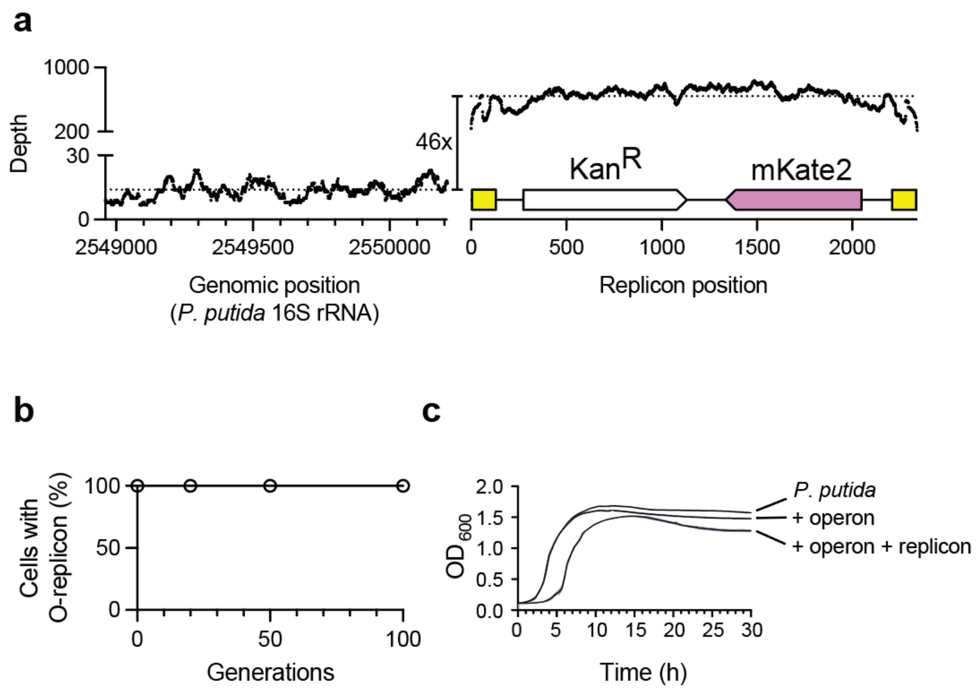


**Supplementary Figure 30. Establishing a synthetic orthogonal replicon in *Pseudomonas putida*.**

**a**, The synthetic replication operon was under the control of an IPTG inducible promoter (PtacIPTG). The $Kan^R$-mKate2 O-replicon consists of flanking 110 bp ITR, a kanamycin resistance gene and an RFP (mkate2) gene. Shown is Illumina sequencing read coverage. **b**, Orthogonal replicons are stably maintained in *P. putida* (n = 3, data are shown as mean± s.d.). **c**, Growth curves of WT *P. putida* and *P. putida* harboring a replicon (n = 8, data are shown as mean ± s.d.).

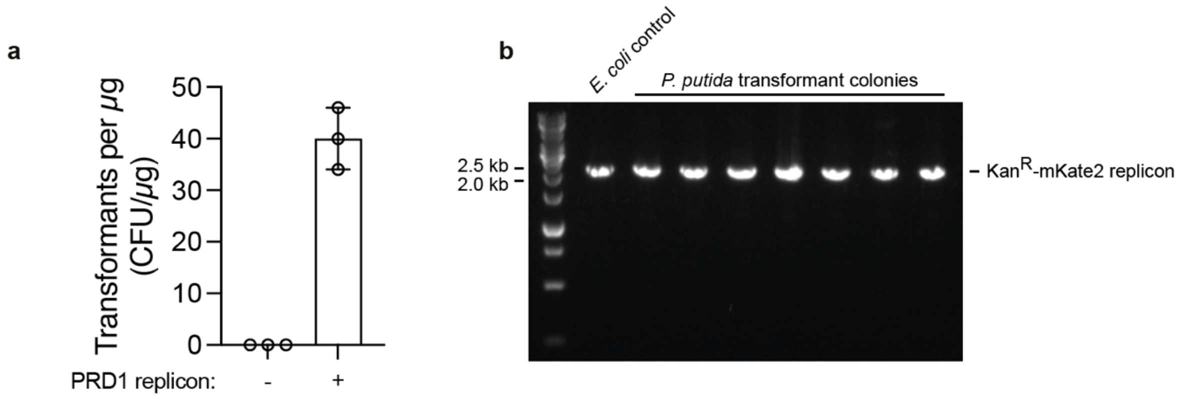


**Supplementary Figure 31. Establishing O-replicons in *P. putida*.** **a**, Efficiency of establishing orthogonal replicons in *P.putida* by electroporating a Kan$^{R}$-mKate2 extracted O-replicon. n = 3, data are shown as mean ± s.d. **b**, Genotyping establishment of the O-replicons in *P. putida*.

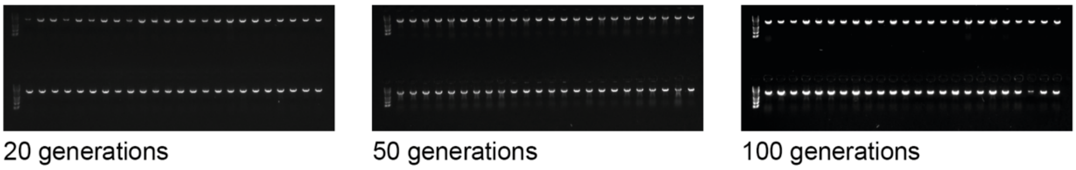


**Supplementary Figure 32. Stability of O-replicons in *P. putida*.** After multiple generations of passaging, 12 colonies were picked from each replicate for genotyping to verify the presence of the O-replicon. All colonies retained the replicon after 100 generations (n = 4).

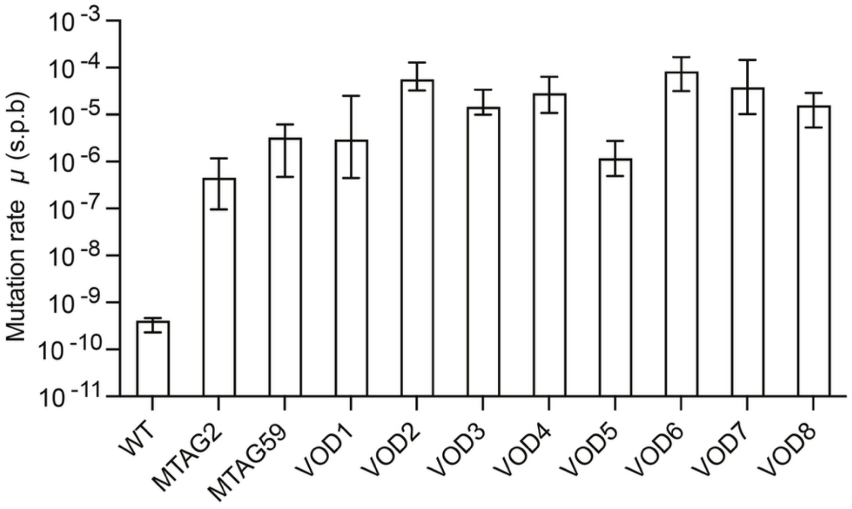


**Supplementary Figure 33. Determination of orthogonal replicon mutation rate (*μ*, s.p.b.) for O-DNAP variants evolved directly in *V. natriegens*.**

The mutation rates were measured after 10 generations via fluctuation tests in *V. natriegens*. For assessment of the orthogonal replicon mutation rate, we used an orthogonal replicon-encoded $Cm^R$ gene with a TAG stop codon at position 38 and the O-DNAP variants were expressed from a p15A plasmid under the control of a constitutive promoter. The identical data for VOD1 and VOD6 are shown in Fig. 3f.

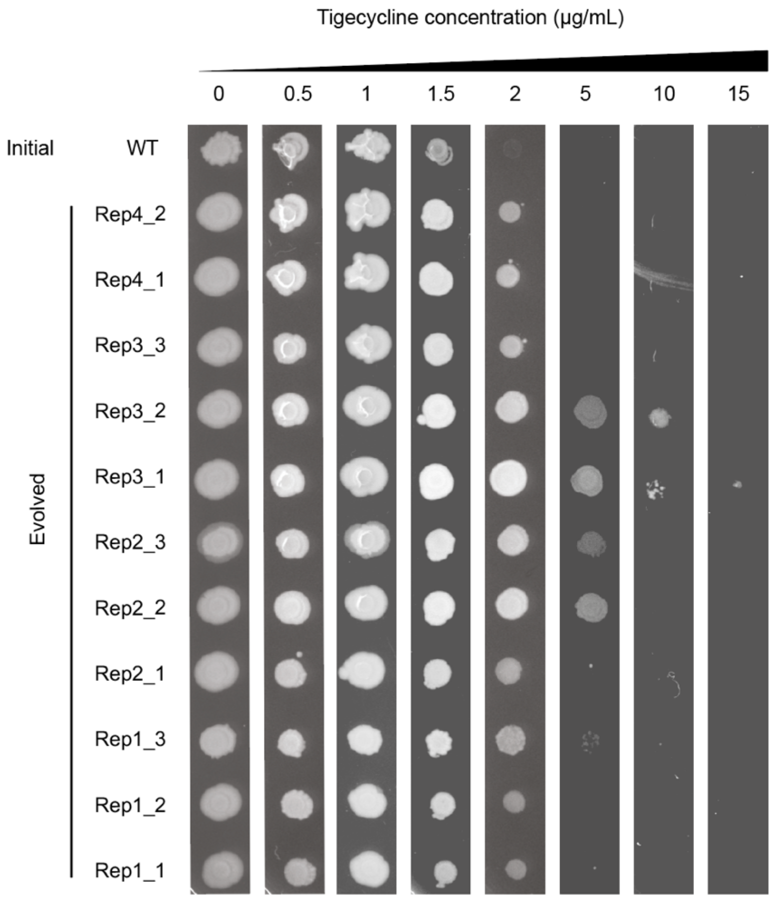


**Supplementary Figure 34. Validation of evolved tetA variants.**

Cells harboring a plasmid (p15A origin) with the evolved *tetA* gene cassette or wild type *tetA* under the control of a PampR promoter were spotted on a range of tigecycline concentrations, as indicated. The identical images for Rep3_2 are shown in Fig. 4d.

| | OrthoRep v1 [1] | OrthoRep v2 [5] | OrthoRep v3 [2] | BacORep v1 [3] | EcORep v1 [6] | EcORep v2 | VinORep v1 |
|---|---|---|---|---|---|---|---|
| Mutation rate (s.p.b.) | $4.00x10^{-8}$ | $1.00x10^{-5}$ | $1.70x10^{-4}$ | $6.80x10^{-7}$ | $9.10x10^{-7}$ | $1.23x10^{-4}$ | $8.18x10^{-5}$ |
| Generations per day | 7 | 7 | 7 | 30 | 28 | 28 | 144 |
| Theoretical mutations per day per kilobase in the absence of selection pressure | 0.00028 | 0.07000 | 1.19000 | 0.02040 | 0.02548 | 3.44400 | 11.77920 |

**Extended Data Table 1. Calculation of theoretical mutations per day per kilobase in the absence of selection pressure across all orthogonal replication systems.**

1. Ravikumar, A., Arrieta, A. & Liu, C. C. An orthogonal DNA replication system in yeast. *Nat. Chem. Bio.* **10**, 175–177 (2014).

2. Rix, G. *et al.* Continuous evolution of user-defined genes at 1 million times the genomic mutation rate. *Science* **386**, eadm9073 (2024).

3. Tian, R. *et al.* Engineered bacterial orthogonal DNA replication system for continuous evolution. *Nat. Chem. Biol.* 1–9 (2023).

4. iGEM Registry of Standard Biological Parts. Anderson Promoter Collection (BBa_J23100–BBa_J23119). http://parts.igem.org/Promoters/Catalog/Anderson.

5. Ravikumar, A., Arzumanyan, G. A., Obadi, M. K. A., Javanpour, A. A. & Liu, C. C. Scalable, Continuous Evolution of Genes at Mutation Rates above Genomic Error Thresholds. *Cell* **175**, 1946-1957.e13 (2018).

6. Tian, R. *et al.* Establishing a synthetic orthogonal replication system enables accelerated evolution in E. coli. *Science* **383**, 421–426 (2024).
